# Mechanistic hierarchical population model identifies latent causes of cell-to-cell variability

**DOI:** 10.1101/171561

**Authors:** Carolin Loos, Katharina Moeller, Fabian Fröhlich, Tim Hucho, Jan Hasenauer

## Abstract

All biological systems exhibit cell-to-cell variability, and this variability often has functional implications. To gain a thorough understanding of biological processes, the latent causes and underlying mechanisms of this variability must be elucidated. Cell populations comprising multiple distinct subpopulations are commonplace in biology, yet no current methods allow the sources of variability between and within individual subpopulations to be identified. This limits the analysis of single-cell data, for example provided by flow cytometry and microscopy. In this study, we present a data-driven modeling framework for the analysis of populations comprising heterogeneous subpopulations. Our approach combines mixture modeling with frameworks for distribution approximation, facilitating the integration of multiple single-cell datasets and the detection of causal differences between and within subpopulations. The computational efficiency of our framework allows hundreds of competing hypotheses to be compared, giving unprecedented depth of a study. We demonstrated the ability of our method to capture multiple levels of heterogeneity in the analyzes of simulated data and data from highly heterogeneous sensory neurons involved in pain initiation. Our approach identified the sources of cell-to-cell variability and revealed mechanisms that underlie the modulation of nerve growth factor-induced Erk1/2 signaling by extracellular scaffolds.

**Highlights:**

- Mechanistic model for populations consisting of multiple heterogeneous subpopulations
- Scalable parameter estimation and model selection using multivariate single-cell data
- Study of pain sensitization in primary sensory neurons
- Proof of principle for fitting of mechanistic model with single-cell data

## Introduction

Cellular heterogeneity is a common phenomenon in biological processes (Elsasser, 1984; De Vargas Roditi and Claassen, 2015). Even isogenic cells of the same cell-type may respond differently to identical stimuli (Tay et al., 2010). This cellular heterogeneity is critical for cellular decision making and the formation of complex organisms (Balázsi et al., 2011). It is also a cause of failure in treatments of cancer, pain, and a wide range of common diseases (Willyard, 2016). Many studies have attempted to gain a deeper understanding of cell-to-cell variability (Rubin, 1990), and recently even a large-scale initiative was found to investigate this heterogeneity (Regev et al., 2017).

Experimentally, most common approaches use methods giving single-cell resolution, such as microscopy (Schroeder, 2011), flow and mass cytometry (Pyne et al., 2009), and single-cell RNA sequencing (Islam et al., 2014). These techniques yield increasing amounts of data, which are commonly analyzed using powerful statistical techniques. Unfortunately, these are unable to identify causalities and latent causes, or to reconstruct the governing equations of the process (see e.g., Moignard et al. (2013)). Improved methods of data analysis are therefore required. We propose a model-based analysis framework for systems exhibiting cell-to-cell variability at different levels:

- differences between cell-types or cellular subpopulations, for example, caused by the cellular microenvironment (Ebinger et al., 2016) or stable epigenetic markers established during cell differentiation (Reik, 2007), and
- differences between cells of the same cell population that arise, for example, from differences in the cell state (Buettner et al., 2015) or from intrinsic stochastic fluctuations (Elowitz et al., 2002).

In the case of homogeneous cell populations, the reaction rate equations (RREs) provide a description of the population behavior in the form of ordinary differential equations (ODEs) (Figure 1A). Stochastic fluctuations or latent differences between cells result in cell-to-cell variability and a distribution of cell states (Hasenauer et al., 2011; Zechner et al., 2012; Yao et al., 2016; Filippi et al., 2016) (Figure 1B). The statistical moments of this distribution are described by moment-closure approximation equations (Engblom, 2006) and system size expansions (van Kampen, 2007; Fröhlich et al., 2016). These methods provide scalable approximations for a range of processes in which variability arises from different sources, the approximation might be cure, e.g. even negative variances might be predicted(Schnoerr et al., 2014). Additionally, they fail to provide an accurate description of the population heterogeneity when subpopulations are present and cannot be used to study the causal differences between cells and subpopulations.

**Figure 1:**
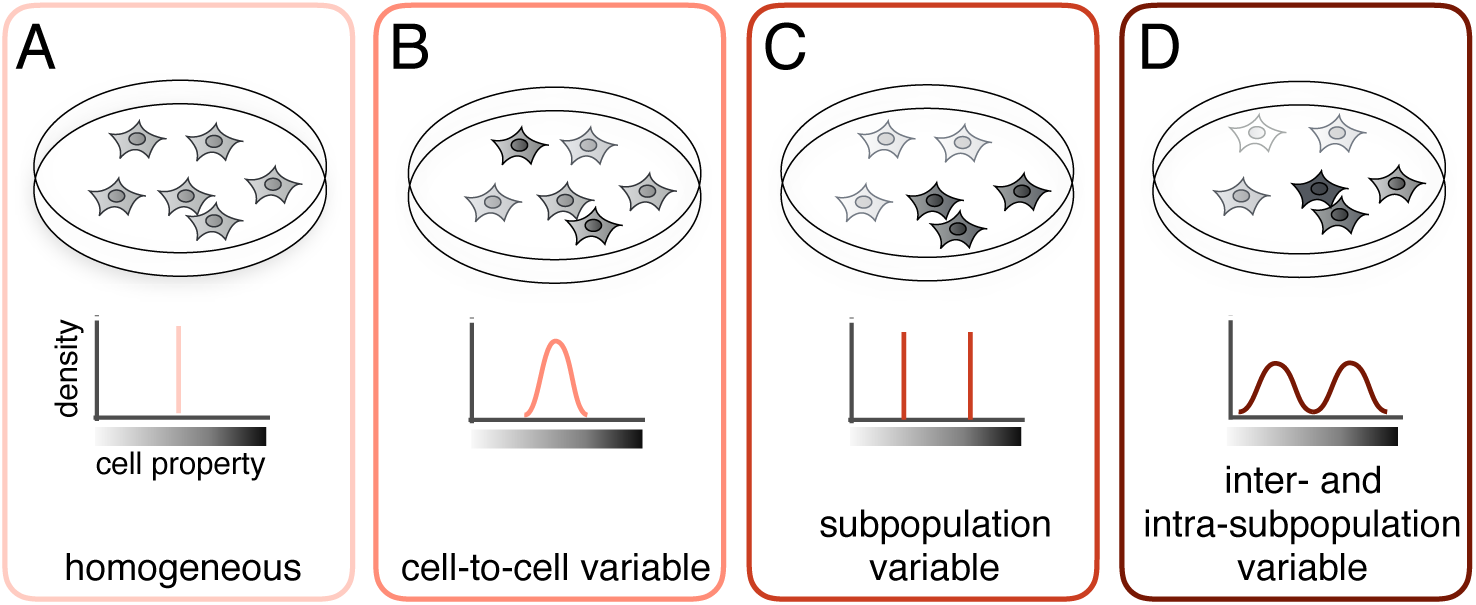
Cell populations exhibiting different levels of heterogeneity. Properties of cells, e.g., receptor levels or reaction rates, indicated by different gray shades for individual cells, can be (*A*) homogeneous: the property is the same for the entire cell population; (*B*) cell-to-cell variable: the property has a unimodal distribution across the cells; (*C*) subpopulation variable: the population can be separated into subpopulations, but within each subpopulation, the property does not vary; (*D*) inter- and intra-subpopulation variable: the property splits the population into subpopulations and also varies between cells within a subpopulation.

To address parameter differences between cell population, Hasenauer et al. (2014) introduced a method that combines mixture modeling and mechanistic RRE modeling of the subpopulation means (Figure 1C). Cell-to-cell variability within a subpopulation is treated naively as an additional parameter that is to be estimated. Thus, the method assumes that the subpopulations are homogeneous and no mechanistic description of cell-to-cell variability within a subpopulation is possible. Moreover, the extant method can only be applied to one-dimensional measurements. When multivariate measurements are used, only marginal distributions can be analyzed and correlations between measurements are neglected, which may result in a substantial loss of information (Altschuler and Wu, 2010; Buchholz et al., 2013).

In this study, we introduced a non-trivial combination of mixture models that is able to capture subpopulation structures and models for individual subpopulations that account for differences between individual cells (Figure 1D). The approach therefore covers several levels of heterogeneity simultaneously (Figure 1A-D). This was not possible using the afore-mentioned approaches, which are all special cases of our model. The means and covariances of the observed species in each subpopulation are linked to a mixture distribution, allowing the entire cell population to be described and providing a mechanistic description of inter- and intra-subpopulation variability. We used the sigma-point approximation (van der Merwe, 2004), a scalable approach allowing for the analysis of large models, to capture the distribution of cell properties within a subpopulation. Similarly, our framework is able to exploit moment equations and system size expansion for the description of individual subpopulations. In contrast with previous work in (Hasenauer et al., 2014), the proposed framework can fully leverage the correlation information in multivariate data, rendering a better conditioned problem and improving identifiability.

We applied this framework to study signal transduction in the extracellular-signal regulated kinase (Erk) pathway, a signaling cascade that is involved in a range of biological processes. Our specific focus was on the pain sensitization of primary sensory neurons in response to nerve growth factor (NGF) stimulation (Hucho and Levine, 2007; Ji et al., 2009; Andres et al., 2012). These neurons encounter a broad range of extracellular environments, including various extracellular scaffolds, and are highly heterogeneous. Performing single-cell microscopy experiments (Andres et al., 2010; Isensee et al., 2014), we investigated the influence of extracellular scaffolds on the response of individual subpopulations. Our computational framework was demonstrated to capture several levels of heterogeneity, including differences between subpopulations as well as differences between cells within a single subpopulation. Our findings suggest that extracellular scaffold structures play a crucial role in pain sensitization signaling and that several underlying mechanistic changes are involved in this.

## Results

### Mechanistic hierarchical population model for single-cell data

We considered populations comprising heterogeneous subpopulations. To allow coverage of multiple levels of heterogeneity, we linked a mixture distribution *ϕ* to a mechanistic model of the means and covariances of individual subpopulations. The distribution of the parameters, e.g., initial conditions or kinetic rates, produces a distribution of cell states and observables (Figure 2A-B). This distribution can be simulated using Monte Carlo methods by drawing parameters from the parameter distribution and simulating the single-cell model. Since this approach is computationally demanding, we approximated the distribution of parameters, states, and observables using finite mixture distributions. The components of the mixture describe the individual subpopulations.

**Figure 2:**
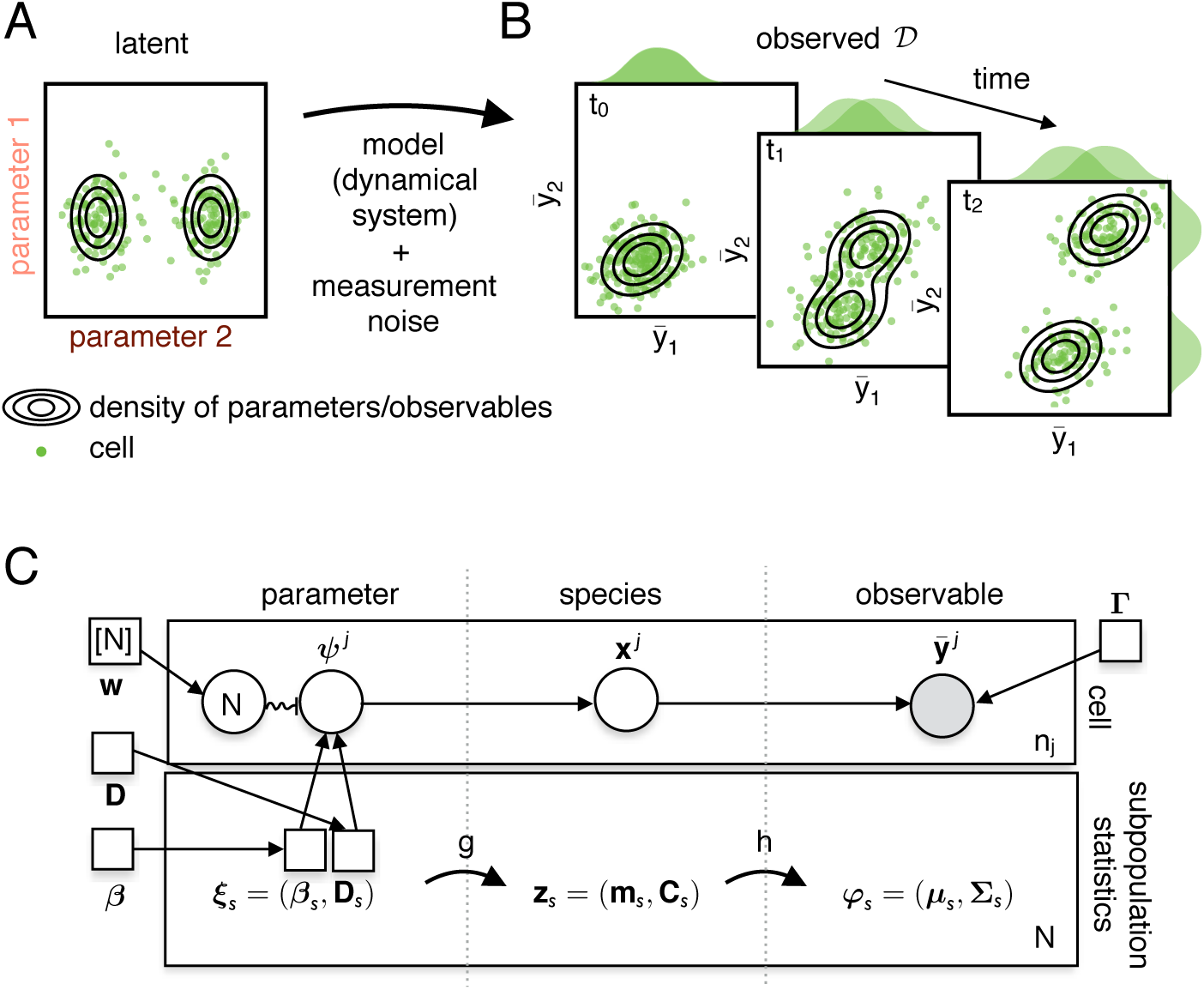
Illustration of the dynamics of a heterogeneous cell population and the mechanistic hierarchical population model. (A) Parameter distribution of a cell population consisting of two subpopulations. The contour lines illustrate the (approximated) parameter density of the cell-to-cell variable parameter 1 and the inter-and intra-subpopulation variable parameters 2. The heterogeneity of parameters is propagated from the latent parameter space to the observed measurement space. (B) Heterogeneity in parameters yields heterogeneous observables **y** = (*y*_1_, *y*_2_)^*T*^ that separate into two subpopulations after stimulation at time point *t*_0_. (C) Structure of the singlecell system and approximation by the hierarchical population model using plate notation. Squares indicate fixed parameters, whereas circles indicate random variables. Gray shading of the circles/squares indicates a known value, whereas the other values are latent. The upper plate illustrates the variables associated with a cell *j*. Each of the *n_j_* cells has parameters *ψ^j^* drawn from a distribution defined by ***ξ**_s_* and **w**. The states of the species **x**^*j*^, resulting from the single-cell dynamics, yield the observables **ȳ**^*j*^, additionally influenced by measurement noise **Γ**. The bottom plate visualizes the statistics of the corresponding cells of a subpopulation. For each subpopulation, the subpopulation parameters ***ξ**_s_* are mapped to the means and covariances of the species of a subpopulation **z**_*s*_, which then are mapped to the distribution parameters *φ_s_*. The observables at the population level are considered to be distributed according to (2).

Each cell *j* has cellular properties encoded in the parameter vector *ψ^j^*. In the hierarchical framework (Figure 2C), these parameters are considered to be drawn from a mixture distribution, as follows:

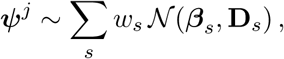

with subpopulation weight *w_s_*, mean ***β**_s_* and covariance **D**_*s*_ for subpopulation *s* = 1,…, *N*. The subpopulation parameters ***ξ**_s_* = (***β**_s_*, ***D**_s_*) classify the variability of a property *ψ_i_* as follows:

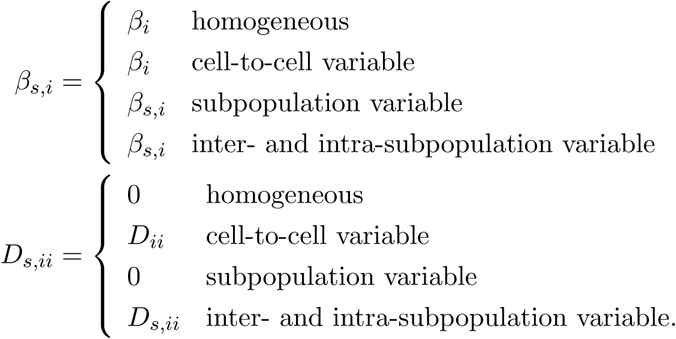

The temporal evolution of the statistical properties of the cells of a subpopulation, including the mean and covariance, are computed using scalable methods. System size expansions and moment equations (van Kampen, 2007; Engblom, 2006) are used to describe stochastic single-cell dynamics, whereas sigmapoints (van der Merwe, 2004) are used otherwise. These approaches yield an ODE model of the statistical moments, comprising the means and covariances **z**_*s*_ = (**m**_*s*_, **C**_*s*_)^*T*^ of species **x**. The model is simulated for each of the *N* subpopulations

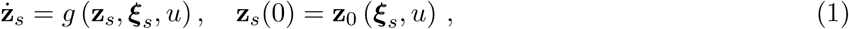

with initial conditions **z**_0_ and experimental condition *u*. The moments of the species in a subpopulation are then mapped to the distribution parameters *φ_s_* = *h* (**z**_*s*_, ***ξ**_s_*, *u*) of the distribution *ϕ*, including measurement noise **Γ**, which is assumed to be the same for all subpopulations. The observables, the quantities of the biological system that can be measured experimentally, are assumed to have the distribution

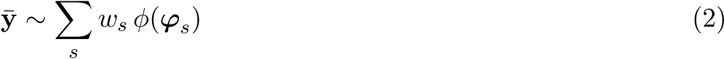

at the population level. In this study, we used mixtures of multivariate log-normal distributions, yielding *φ_s_* = (***μ**_s_*, **Σ**_*s*_). The sigma-point approximation (detailed in supplementary Information) provides time-dependent moments of the system defined in (1) and accounts for cell-to-cell variability. When combined with subpopulation variability, this yields both the inter- and intra-subpopulation variability. For a comparison of our approach to existing methods, we refer to the supplementary Information.

### Parameter estimation and model selection

The parameters of biochemical processes, the sources of cell-to-cell and subpopulation variability, and the precise network structure are in general unknown. We therefore calibrated the hierarchical population model using single-cell snapshot data **ȳ**^*e*,*k*,*j*^ with cell *j* measured at time point *t_k_* under experimental condition *u_e_*, for example, representing a drug dosage. The parameters ***θ*** ∈ ℝ^*n_θ_*^ usually comprise characteristics of a subpopulation (e.g., the means and covariances of the parameter distributions), subpopulation sizes and measurement noise. Maximum likelihood estimation was used to derive these parameters from the data. The maximum likelihood estimate ***θ̂*** was obtained by solving the following optimization problem:

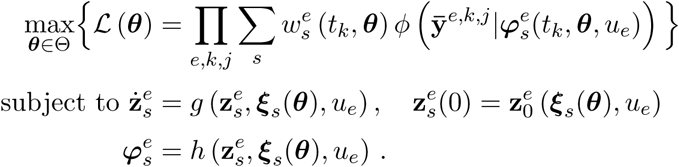

The likelihood function 𝓛 incorporates all cells, time points, and experimental conditions. For efficient parameter estimation, we performed multi-start local optimization with a robust evaluation scheme for the log-likelihood function and its gradient. The gradient of the log-likelihood function with respect to the parameters was computed using forward sensitivity analysis (see (Raue et al., 2009; Loos et al., 2016) and supplementary Information).

To infer the subpopulation structure, the difference between subpopulations, the variability within subpopulations, and the influence of the experimental condition, a collection of hierarchical models is formulated. We compare these models and the corresponding hypotheses using the Bayesian Information Criterion (BIC) (Raftery, 1999).

The hierarchical models were implemented in the MATLAB toolbox, incorporating efficient simulations for the individual subpopulations. While any simulation that provides means and covariances of the subpopulations can be employed, in this study, we used the sigma-point approximation. This approach accounts for cell-to-cell variability, which is manifested in the parameters (see supplementary Information for more details).

### Unraveling sources of heterogeneity

To demonstrate the advantages of the hierarchical population model, which incorporates a mechanistic description of the means and variances, over the method proposed by Hasenauer et al. (2014), we applied our approach to simulated data on a simple conversion process. Such conversions are common in biological systems, for example, in phosphorylation. The conversion process comprised two species A and B, with cell-to-cell variable conversions from B to A (Figure 3A), corresponding to different levels of phosphatase in the cells. Two subpopulations were assumed with different responses to stimulus *u*. This produced subpopulations with different rates of stimulus-dependent conversion from A to B. Artificial measurement noise was added to allow the capability of the framework to distinguish measurement noise from biological variability to be assessed. We assumed the underlying subpopulation structure, i.e., the subpopulation variability of *k*_1_, to be known (detailed in supplementary Information).

**Figure 3:**
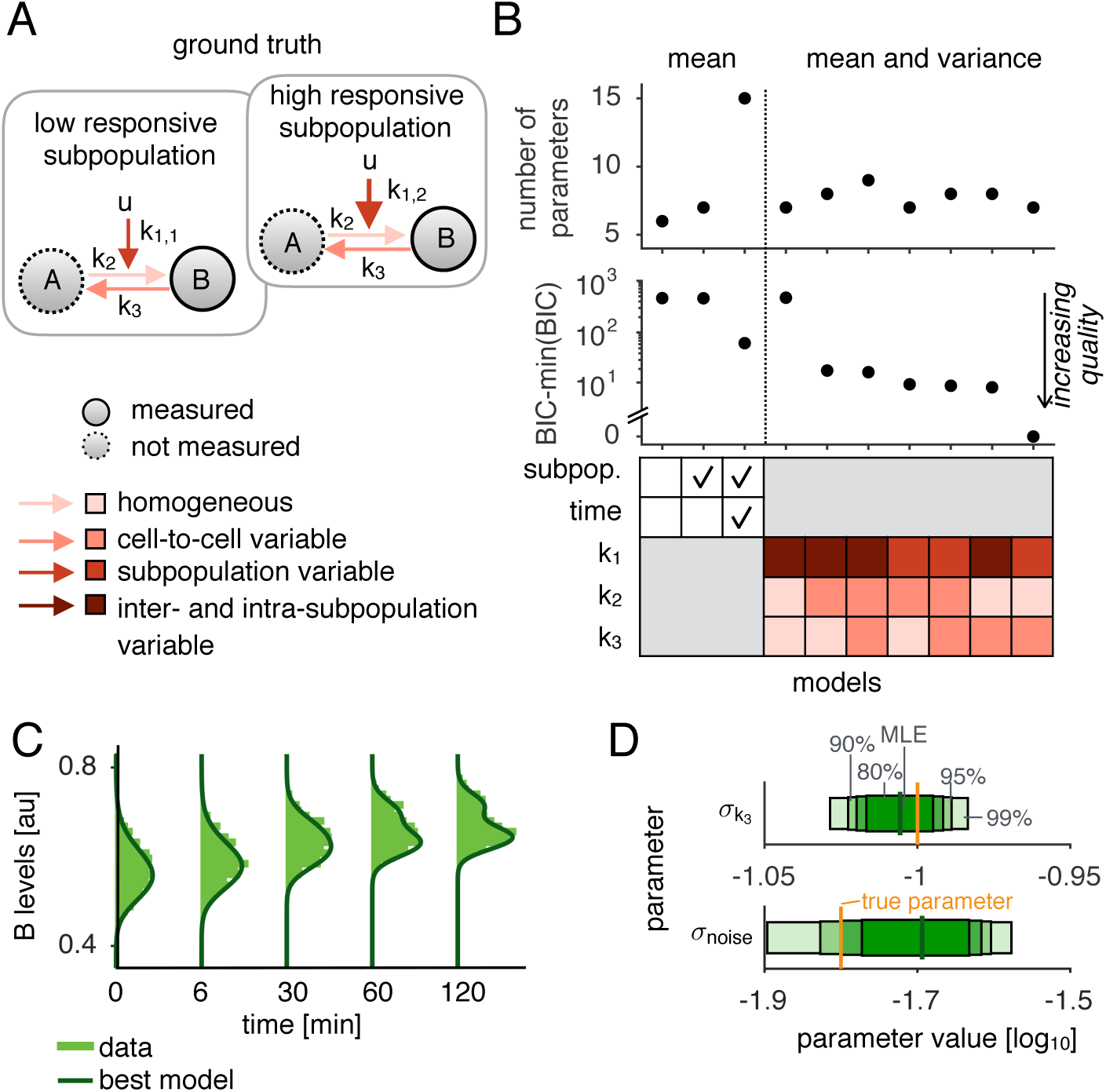
Inference of cell-to-cell variability using mechanistic models. (A) Model of a conversion between two species *A* and *B* comprising two subpopulations differing in their response to stimulus *u*. Different colors indicate the variability of the reaction rates. (B) Model selection with the Bayesian Information Criterion (BIC). The first three models use RREs according to (Hasenauer et al., 2014) and vary in the number of additional parameters (1, 2, and 10) for the variances of the mixture distribution. The last models use the mean and variance obtained by sigma-points and differ in their sources of heterogeneity. (C) Data on the conversion process (1000 cells per time point) and fit corresponding to the best and true underlying model. (D) Confidence intervals for the variability of *k*_3_ and the measurement noise (*σ*_noise_). Horizontal bars show the confidence intervals corresponding to the 80%, 90%, 95%, and 99% confidence levels, and the vertical lines the maximum likelihood estimates (MLE).

The simulated data were analyzed using (i) the approach introduced by Hasenauer et al. (2014) which describes the subpopulations using RREs and (ii) the proposed approach using hierarchical single-cell analysis. The first approach does not model the temporal evolution of the variance, requiring different parameterizations to be compared, i.e., constant, time-dependent, and time/subpopulation-dependent variability. Model selection with the BIC indicates that different parameters for each subpopulation at every time point are required to be used to describe the data (Figures 3B and S1). This demonstrates that the observed cell-to-cell variability changes over time but provides no information about the sources of the observed cell-to-cell variability.

The mechanistic modeling of multiple levels of heterogeneity facilitates the identification of its causal source via model selection. We considered a range of hypotheses and performed model selection using BIC (Figure 3B). Given the subpopulation structure, the additional source of heterogeneity, namely, the conversion from B to A, was correctly identified using the BIC and the corresponding model provided a good fit to the data (Figure 3C). The BICs for most of the hierarchical models were substantially lower than that of the best model that incorporates only the mean. This confirms that a mechanistic description of the variability is more appropriate.

We analyzed the ability of the hierarchical model to identify the different contributions of cell-to-cell variability and measurement noise, as both are normally present in single-cell experiments. The uncertainty analysis suggested that the hierarchical modeling approach was able to distinguish between the two (Figure 3D).

This example shows how the hierarchical population model outperforms the variants of models presented by Hasenauer et al. (2014). We confirmed the power of the proposed approach by studying a model of stochastic gene expression (Figure S6) and comparing the approach to the method by Zechner et al. (2012) (see supplementary Information). Our model employs a mechanistic description of the variability, thereby enabling a more detailed insight into the heterogeneity of the population and reducing the number of parameters that need to be estimated from the data.

### Identification of differential protein expression using multivariate data

Many single-cell technologies provide multivariate measurements and therefore convey information about the correlations between the observables. To incorporate this, we extended our hierarchical modeling framework to multivariate data and demonstrated its capability to reconstruct the differential protein expression of cellular subpopulations (Sauvageau et al., 1994; Kharchenko et al., 2014) using simulated data. We considered a model describing the abundance of two proteins, the expression of which is regulated by stimulus *u* (Figure 4A). The influence of *u* varies between cell populations and is therefore able to capture, e.g., different levels of membrane receptors. We generated multivariate data by simulating a single-cell model (see supplementary Information for more details).

**Figure 4:**
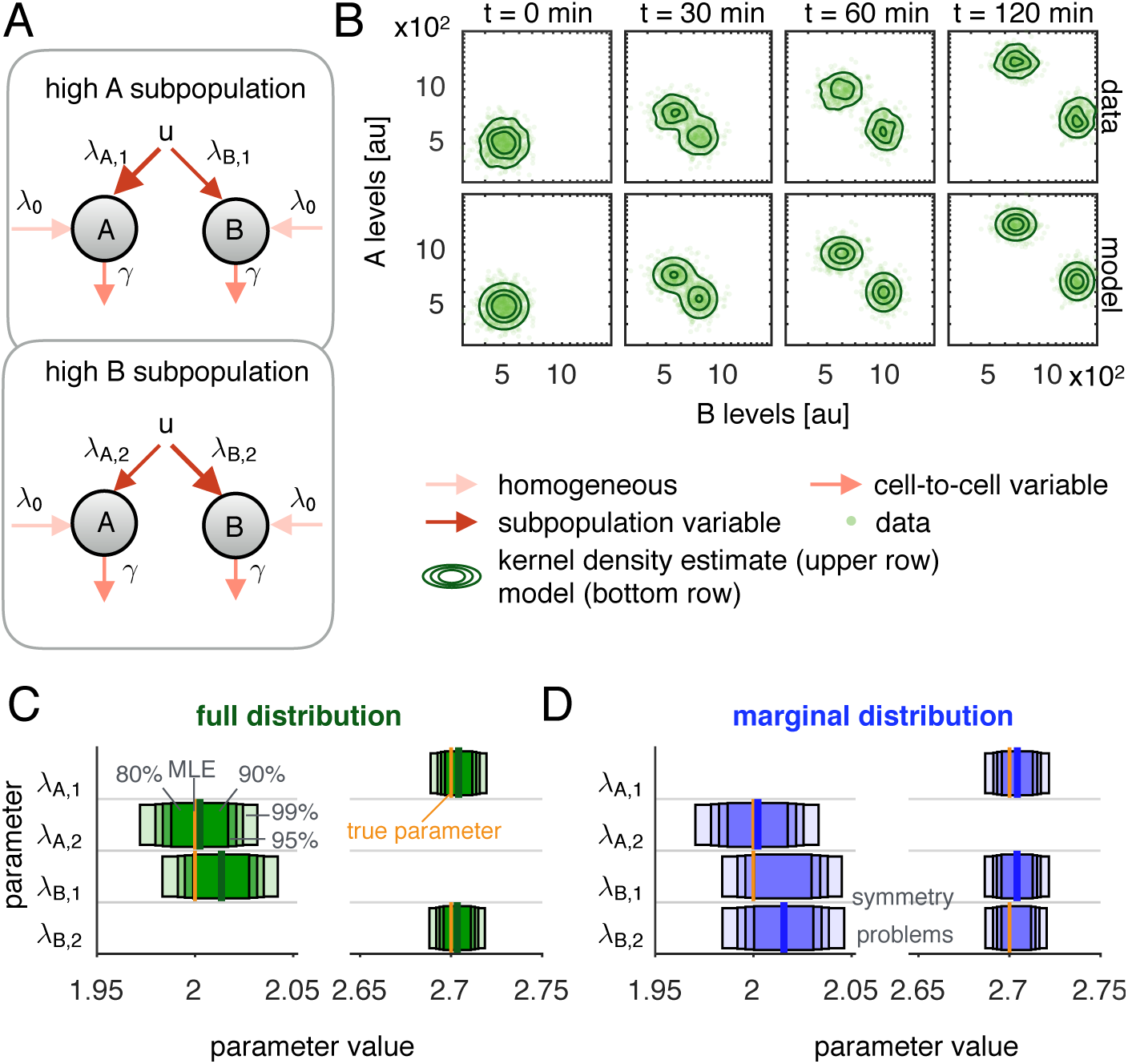
Reconstruction of differential protein expression in heterogeneous populations using multivariate data. (A) Model of differentially expressed proteins A and B. (B) Upper row: data points (1000 cells per time point) and kernel density estimation. Lower row: data points and model for the full distribution. (C,D) Confidence intervals for the parameters of the model using (C) the full distribution and (D) the marginal distributions. Horizontal bars show the confidence intervals corresponding to the 80%, 90%, 95%, and 99% confidence levels. The vertical lines show the MLE.

An analysis using our hierarchical approach confirmed the ability of the proposed model to reproduce the data (Figure 4B) and to provide reliable parameter estimates (Figure 4C). Such multivariate data cannot be exploited by the existing model-based approaches. When the temporal evolution of proteins is measured individually, the correlation information is missing and a symmetry arises in the system (Figure 4D). This is reflected in the multimodal profiles of the parameters λ_B,1_ and λ_B,2_, indicating a lack of practical identifiability.

Our framework exploits the correlation structures of multivariate data, which in this simulation example allowed us to conclude that each subpopulation had a high expression of only a single protein. This only becomes possible when the correlations are analyzed.

### Modeling signal transduction in neuronal populations

We applied the hierarchical modeling approach to investigate the influence of an extracellular scaffold on NGF-induced Erk1/2 activation in cultures of adult sensory neurons (Figure 5A). This was done by monitoring the rates of NGF-mediated Erk1/2 phosphorylation in dissociated cultures of the primary sensory neurons of rat dorsal root ganglia. NGF-mediated Erk1/2 signaling has been shown to play a crucial role in nociceptor sensitization in thermal and mechanical hyperalgesia (Zhuang et al., 2004; Malik-Hall et al., 2005). Primary sensory neurons form a heterogeneous population, from which, upon NGF stimulation, a subpopulation reacts with a graded Erk1/2 phosphorylation response. Previous models have attempted to approximate this by assuming the existence of responders and non-responders with differing levels of the NGF receptor TrkA (Hasenauer et al., 2014). In the current study, we refined this substantially by modeling the overall population using two heterogeneous subpopulations that differed in their average response. To calibrate this refined model, we collected quantitative single-cell microscopy data on NGF-induced Erk1/2 phosphorylation kinetics and dose response curves using immunofluorescence labeling of pErk1/2 alone, co-labeled with Erk1/2 and TrkA antibodies (see supplementary Information for more details).

Our analysis used the ODE model introduced in (Hasenauer et al., 2014). This has six structurally identifiable parameters *k*_1_, *k*_2_, *k*_4_, *k*_5_, *k*_3_[TrkA]_0_ and *c*[Erk]_0_. Based on the experimental observations, we assumed cell-to-cell variability in relative Erk1/2 expression and in cellular TrkA activity.

### Causal differences between neuron subpopulations

To study the differences between responders and non-responders, we used experimental kinetic and dose response data from sensory neurons cultured on the adherence substrate poly-D-lysine (PDL). We fitted 64 models accounting for all combinations of the six potential differences between subpopulations, which was only feasible due to the computational efficiency of our approach. Our assessment of the importance of individual differences between the subpopulations using a BIC-based ranking scheme suggested that cellular TrkA activity (*k*_3_[TrkA]_0_) made the greatest contribution (Figure 5B). This was indicated by a high BIC weight, which captures differences by Bayesian model averaging (see supplementary Information for more details), and the substantially better mean rank of the models using differences in cellular TrkA activity compared with those using other differences. The additional subpopulation variability of TrkA expression levels was also confirmed experimentally in the cultures (Figure 5D) and use of this difference alone produced an excellent fit to the experimental data (Figures 5C and S4). The following potential differences are the relative Erk1/2 expression levels (*c*[Erk]_0_) and the dephosphorylation rate (*k*_5_). However, our experimental data showed no statistically significant difference in total Erk1/2 levels between responders and non-responders (Figure 5E). To assess the relevance of the dephosphorylation rate and thus the corresponding phosphatase activity we performed experiments in which we monitored the pErk1/2 decline dynamics after inhibiting the mitogen-activated protein kinase (Mek) that phosphorylates Erk1/2. If the phosphatase activity does vary, we would expect to observe different equilibration dynamics. However, this could not be confirmed (Figures 5F and S3). This demonstrated that the hierarchical approach provided an appropriate ranking of differences and suggested that the total TrkA levels differed between subpopulations.

**Figure 5:**
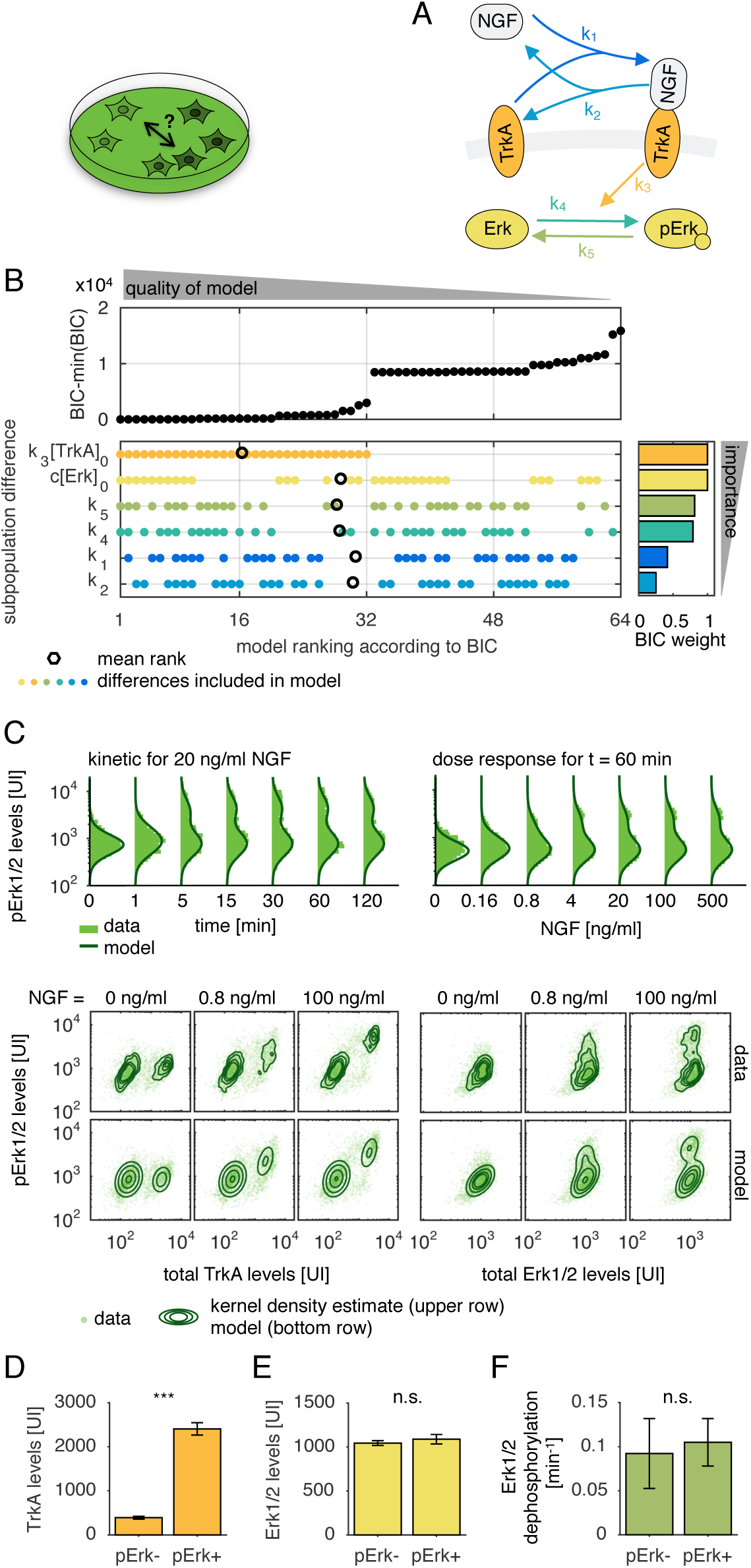
Detecting the sources of heterogeneity between subpopulations in primary sensory neurons. (A) Pathway model of NGF-induced Erk signaling. (B) Ranking according to the BIC values for the 64 hierarchical models, in which the colored dots indicate those parameters that are assumed to differ between the subpopulations. The importance of the differences is ranked according to the BIC weights. The black circles indicate the mean rank of the models including the corresponding difference. (C) Data and fit for measurements of pErk1/2 levels (approximately 1400 cells per time point and 4300 cells per dosage) and multivariate measurements of pErk/TrkA and pErk/Erk levels (approximately 3000 cells per dosage) measured for 60 min under NGF stimulation with indicated concentrations. The measured values are in arbitrary units of intensity. For the multivariate data, the contour lines of the kernel density estimation of the data and the level sets of the density of the hierarchical model are shown. Mean and standard deviation of (D) TrkA levels (*n_r_* = 4 replicates) (E) Erk1/2 levels (*n_r_* = 4) and (F) Erk1/2 dephosphorylation (*n_r_* = 4) of non-responsive (pErk-) and responsive (pErk+) sensory neurons after NGF stimulation with varying concentrations (as indicated in (C) for 60 min).

### Influence of extracellular scaffolds on sensitization signaling

It is well established that during painful conditions, a plethora of different soluble molecules in the extracellular space, including cytokines, neurotrophins, and neuropeptides, induce nociceptor sensitization (Hucho and Levine, 2007). It is also known that the extracellular scaffolding structures, namely extracellular matrix molecules, are often highly altered in painful conditions such as wounds or tumors. However, much less is known about the role of cell scaffolds in sensitization of nociceptive neurons. We therefore compared the modification of NGF-induced sensitization signaling of sensory neurons by collagen type I (Col I), a classical extracellular matrix protein that forms receptor-matrix interactions, and by poly-D-lysine (PDL), a neutral scaffolding that promotes cell adherence by electrostatic interaction. We determined the kinetics and dose response curves of NGF-induced Erk1/2 phosphorylation in sensory neurons cultured overnight on Col I or PDL (see supplementary Information for more details). It was found that the mean Erk1/2 activation was approximately 17% higher in Col I compared to PDL after NGF treatment (Figure 6A for pErk1/2 dose responses and Figure S5A for the other datasets). In addition to showing increased NGF-induced Erk1/2 activation, the number of cells was observed to be 1.5 times lower in the collagen cultures than in the polylysine cultures. These observations raised questions about the source of the measured increase in mean NGF-mediated Erk1/2 activation. We considered two hypotheses: (i) the increase results from a biological action of the different scaffolds onto the neurons and (ii) the increase reflects a shift of the subpopulation sizes arising from a nonrandom loss of parts of the high-responder subpopulation due to reduced cell adherence in the collagen cultures. To unravel the causal differences between the primary sensory neurons cultured on PDL and on Col I, we applied 128 hierarchical models using the previously derived subpopulation structure. These models considered all combinations of differences between the cell population on different scaffolds, including the size of subpopulations. The model for each adherence substrate accounted for the cell-to-cell variability of Erk1/2 and the inter- and intra-subpopulation variability of cellular TrkA activity. The model ranked first by the BIC (Figures 6B) gave a good fit to the data and took account of differences in Erk1/2 expression (*c*[Erk]_0_), Erk1/2 dephoshorylation (*k*_5_), and cellular TrkA activity (*k*_3_[TrkA]_0_) (Figures 6C-D and S5B-D). These differences were assumed to explain the higher response on Col I, and therefore supported hypothesis (i). The model that assumed no difference between the extracellular scaffolds (rank 128) or changes only in the relative size of the subpopulations (rank 127) performed worst, indicating that hypothesis (ii) failed to explain the data. The differences in relative TrkA and Erk1/2 expression levels were also observed experimentally (Figures 6D-E). These results confirmed the model-based analysis and suggested an impact of the classical extracellular matrix protein collagen I on protein expression.

**Figure 6:**
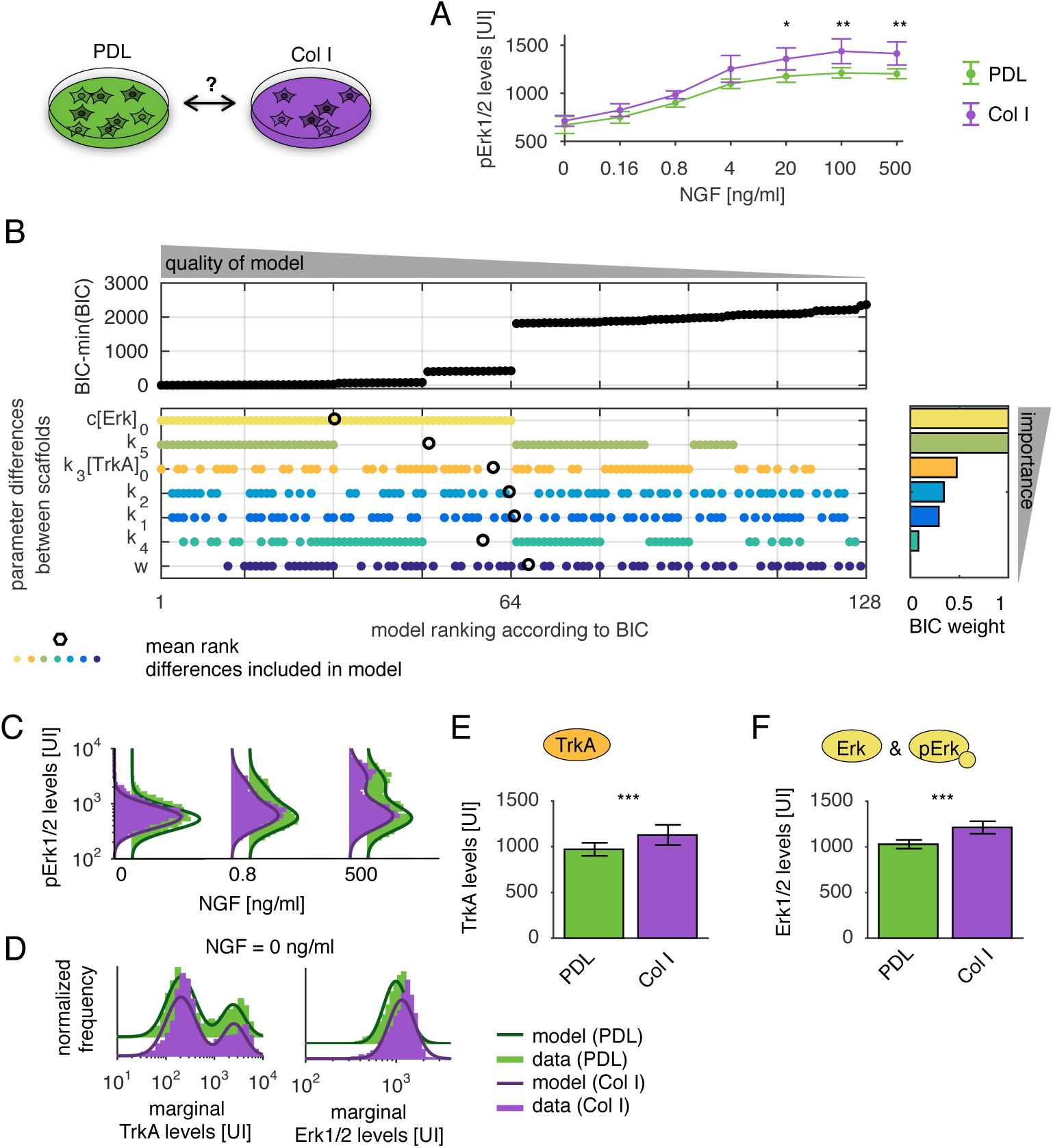
Differences in NGF-induced Erk1/2 phosphorylation mediated by different extracellular scaffolds. Primary sensory neurons were provided with the two different scaffolds poly-D-lysine (PDL) and collagen type I (Col I) in an overnight culture. (A) Sensory neurons grown on the Col I substrate showed a significantly higher mean phospho-Erk1/2 response to indicated doses of NGF after 1h of stimulation. Means and standard deviations of four replicates are shown. (B) BIC-based ranking for the potential differences between culture conditions. The colored dots indicate which parameters are assumed to differ between the extracellular scaffolds. (C) Experimental data and fit for measurements of pErk1/2 distributions from Col I (approximately 2300 cells per dosage) and PDL (approximately 4300 cells per dosage) cultured neurons after treatment with indicated NGF concentrations for 1 h. (D) Marginal levels for TrkA and Erk1/2, which were assumed to be constant over varying doses and time (approximately 2000 cells in Col I and 2900 in PDL). Mean and standard deviation of (E) TrkA and (F) Erk1/2 levels of NGF dose response curve data, which showed significant elevations in Col I treated neurons. For this calculation, 24 samples were used (4 replicates for 6 doses).

## Discussion

Elucidating the causes of cellular heterogeneity is a challenging task in systems biology and requires appropriate mechanistic models for use with single-cell data. In this study, we proposed a hierarchical modeling framework that for the first time allowed different levels of heterogeneity to be investigated, including subpopulation structures and cell-to-cell variability within subpopulations. It also provides mechanistic insights. Beyond cell-to-cell variability, the method accounts for measurement noise and is able to deconvolute these sources.

This modeling approach unifies available description frameworks (Zechner et al., 2012; Hasenauer et al., 2014) and exploits efficient simulation methods for cellular subpopulations. We focused on the cell-to-cell variability encoded in parameter values (Koeppl et al., 2012) and used sigma-point approximation to determine the subpopulation means and variances. To address variability arising from stochastic fluctuations, moment equations (Figure S6) and other methods, including the system size expansion (Fröhlich et al., 2016), can be used. The proposed method facilitates the integration and simultaneous analysis of multiple datasets, without requiring complex pre-processing of the data (Lee et al., 2011). The modeling approach is implemented in the open-source MATLAB Toolbox ODEMM which is available on GitHub and ready to be reused by the community.

Differences between cell types can be analyzed and modeled in the same manner as differences between cellular subgroups. The method is also able to handle more than two subpopulations, and the number of subpopulations might even be inferred using a data-driven approach.

To demonstrate and evaluate the method, we collected a comprehensive dataset on NGF-induced Erk1/2 signaling in primary sensory neurons, a challenging experimental system. We investigated the subpopulation structures and assessed the influence of different scaffolding environments on the signaling pathway and the composition of subpopulations. Using simultanous fitting of dose responses and activation kinetics for the different adherence substrates, our method revealed an association between collagen type I and elevated NGF-mediated Erk1/2 signaling, demonstrating that extracellular scaffolds play a role in nociceptor sensitization signaling. In this study, we compared Col I with the artificial PDL substrate, but a wide range of extracellular matrix proteins are altered in conditions such as wounds and tumors. Further analysis will be needed to determine the impact of single extracellular matrix components on sensitization signaling of nociceptive neurons. Further research is also needed to investigate whether modulated signaling leads to increased excitability of these neurons.

Procedures such as a forward-backward algorithm (e.g., Hastie et al. (2009)) or reversible jump Markov Chain Monte Carlo (Green, 1995) could be implemented to simultaneously perform parameter estimation and model selection. In this study, mixtures of log-normal distributions were used to model the cell population. However, other distributions, including the Laplace distribution, could be integrated with the computational framework to improve robustness against outliers (Maier et al., 2017).

The inference of mechanistic models from single-cell data relies on statistical models for the measurement and sampling process. In many modeling studies using single-cell data, no distinction is made between cells from different batches, obscuring cell-to-cell variability and differences between experimental batches (Hicks et al., 2015). In this study, we observed that the derived likelihood function can be overly sensitive and that model selection is biased towards complex models. To circumvent this issue, we used a ranking of potential differences rather than a precise measure of statistical significance. However, this problem will need to be addressed, as the use of single-cell data is increasingly common.

In summary, we proposed the use of hierarchical population models as a novel tool to study heterogeneity in multivariate single-cell data and evaluated their performance. Our framework is the first to account for multiple levels of heterogeneity simultaneously. Our results on simulation and application examples suggest that this method can be used to obtain a more holistic understanding of heterogeneity.

## Author Contributions

C.L., F.F., and J.H. developed the method. K.M. and T.H. designed the experiments. K.M. performed the experiments. C.L. analyzed the data. C.L., K.M, F.F., T.H., and J.H. wrote the paper.

## Acknowledgements

C.L., F.F., and J.H. acknowledge financial support from the German Research Foundation (DFG) and the Postdoctoral Fellowship Program of the Helmholtz Zentrum München. K.M. was supported by the Evangelisches Studienwerk Villigst and the graduate program in Pharmacology and Experimental Therapeutics at the University of Cologne which is financially and scientifically supported by Bayer. This work was supported by the German Research Foundation (DFG; http://www.dfg.de) through the Graduate School of Quantitative Biosciences Munich (QBM; FF).

## Supplementary Information

### Experimental model

#### Antibodies

The following antibodies were used in this study: chicken polyclonal antibody against UCHL1 (1:4000; Novus, #NB110-58872), mouse monoclonal antibody against UCHL1 (1:1000, MorphoSys, #7863-2004), rabbit monoclonal antibody against phospho-Erk1/2 (1:250, Cell Signaling, #4370L), mouse monoclonal antibody against ERK1/2 (1:500, Cell Signaling, cat# 4696 S), goat polyclonal antibody against TrkA (1:500, R&D Systems, #AF1056), and highly cross adsorbed Alexa Fluor 488, Alexa Fluor 568, Alexa Fluor 594-, and Alexa Fluor 488-conjugated secondary antibodies (Invitrogen).

#### Reagents

NGF (50*μ*g/ml in 0.1 % BSA), GDNF (20*μ*g/ml in 0.1 % BSA), U0126 (50 mM in DMSO) were purchased from Alomone labs (#N-240), PeproTech (cat# 450-51), and Calbiochem (# 662005), respectively, and were prepared as indicated. Concentrations used are indicated in the text or figure legend. Collagen type I (Cell Systems, #5056-A) and poly-D-lysine (Sigma,# P6407-5MG) were diluted in 1xPBS to final concentrations of 3.4 *μ*g/ml and 10 *μ*g/ml.

#### Animals

Male Sprague Dawley rats (200250 g, 8-10 weeks old) were obtained from Harlan Laboratories. All experiments were performed in accordance with the German animal welfare law with permission of the District Government for Nature and Environment, NRW (LANUV NRW, license 84-02.05.20.13.045). Rats were sacrificed by CO_2_ intoxication for tissue isolation.

#### Coating

96-well imaging plates (Greiner) were coated with 50 *μ*l volume of matrix protein dilutions per well for 3h at 37 C. Wells were washed one time with 1xPBS for 10 min. PDL coatings were dried and washing solution of Col I treated wells was removed immediately before cell seeding.

### Primary sensory neuron culture

L1-L6 dorsal root ganglia (DRG) were isolated, desheathed, pooled and incubated in Neurobasal-A (NB) medium supplemented with collagenase P for 1 h in 5 % CO_2_ atmosphere at 37 C. Neurons were dissociated by trituration with fire-polished siliconated Pasteur pipettes and axonal debris and disrupted cells were removed by a 14% BSA gradient centrifugation (120g, 8min). Cells were resuspended in NB medium supplemented with B27 medium, L-Glutamine, L-Glutamate and Penicillin-Streptomycin. subsequently, they were plated on pre-coated 96-well imaging plates and incubated overnight in a 5 % CO_2_ atmosphere at 37 C.

### Stimulation and fixation of neuronal cultures

Neuronal cultures were stimulated 15 h after isolation by removal of 50 *μ*l culture medium, mixing with the compound and returning to the corresponding culture well. Solvent controls were treated alike. Stimulation was performed with automated eight-channel pipettes (Eppendorf) on pre-warmed heating blocks (37 C), and stimulated cells were placed back into the incubator. Neurons were fixed by adding 8 % PFA (final concentration 4% PFA) for 10 min at RT and subsequently washed three times with 1xPBS for 10min. Kinetic experiments involved time courses of 0, 1, 5, 15, 30, 60 and 120 min NGF stimulation (20ng/ml), whereas dose response curves were obtained by NGF stimulations with the following NGF concentrations for 1 h: 0.16, 0.8, 4, 20, 100, 500ng/ml.

### Immunocytochemistry

Cells were blocked and permeabilized with 2% normal goat serum or 2 % normal donkey serum supplemented with 1 % BSA, 0.1 % Triton X-100, 0.05% Tween 20 for 1 h at RT. Primary antibodies were added in 1 % BSA in 1xPBS and cells were incubated overnight at 4 C. After three washes with 1xPBS for 10 min at RT, cells were incubated with secondary antibodies diluted in 1xPBS for 1 h at RT. Plates were stored at 4 C after three additional washes with 1xPBS (10 min, RT) until scanning.

### Quantitative microscopy

Immunofluorescently labelled neurons were imaged via the Cellomics ArrayScan microscope using a 10x objective as described previously (Isensee et al., 2014). Images of 512 × 512 pixels were analyzed using the Cellomics software package. Briefly, images of all channels were background corrected (low pass filter), objects were identified using fixed thresholding (intensity 900) and segmentation by shape (parameter 15). Neurons were validated by the following object selection parameters: size: 1657500*μ*m^2^; circularity (perimeter^2^/4*π* area): 12; length-to-width ratio: 12.67; average intensity: 90012.000; total intensity: 2 10^5^ to 5 10^7^. The image masks were then used to quantify signals in other channels. Raw values of three to four independent experiments were further processed via the R software. Raw fluorescence data was compensated and normalized. In brief, three controls were prepared for a triple staining: 1. UCHL1 alone, 2. UCHL1 + antibody 1, and 3. UCHL1 + antibody 2. Raw fluorescence data of the controls were used to calculate the bleed-through between fluorescence channels. The slope of best fit straight lines were determined by linear regression and used to compensate bleed through as described previously (Roederer, 2002). Compensated data were scaled to a mean value of 1000 for the unstimulated cells of the poly-D-lysine control to adjust for variability between experimental days.

## Methods

### Models for individual subpopulations

The hierarchical modeling approach introduced in this manuscript describes the population dynamics based on the dynamics of individual subpopulations. In this section, we introduce the modeling approaches at the subpopulation level that are used in our study.

First, we considered the simple case that only the mean of a subpopulation is modeled mechanistically, whereas the variance and higher order moments are not linked to the underlying biochemical reaction network. For this, the reaction rate equation (RRE) was used

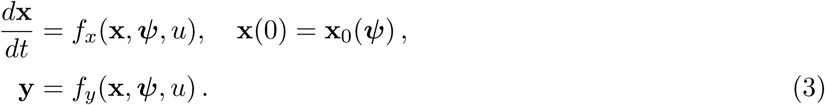

Here, **x** = (*x*_1_,…, *x_n_*) denotes the biochemical species, **y** = (*y*_1_,…, *y_d_*) the observables of the system, and *ψ* the parameters, such as reaction rates, protein abundances, or initial conditions. This follows the method introduced by (Hasenauer et al., 2014).

The RRE is based on the assumption that the subpopulations are homogeneous. However, many cellular processes exhibit substantial intrinsic or extrinsic cell-to-cell variability. To account for this variability, we considered models accounting for random parameters and stochastic reaction kinetics.

#### Sigma-point approximation

In this study, we modeled extrinsic variability by heterogeneity in *L* parameters of the parameter vector *ψ* ∈ ℝ^*n_ψ_*^ of individual cells. The parameters *ψ* were assumed to follow a probability distribution *p_ψ_* (*ψ*). This distribution in the parameters *p_ψ_* (*ψ*) is mapped to a distribution of cell states and observables of the subpopulation, which need to be computed for the parameter estimation. A detailed analysis of this image requires sampling from *p_ψ_* (*ψ*) and subsequent evaluation of the state and observable vectors by simulation. This procedure is, however, computationally demanding. We employed the sigma-point approximation (van der Merwe, 2004) to obtain an approximation of the statistical moments of the image, mean and covariance and their dynamics in time, using a small number of simulations. The sigma-point approximation uses only the image of 2*L* + 1 deterministically chosen parameter vectors. These parameter vectors, the so called sigma-points, are chosen to represent the mean *β* and the covariance **D** of *p_ψ_*. For the parameters that were considered to be homogeneous, i.e., not variable across the cells, it was assumed that *β_i_* = *ψ_i_* and *D_ii_* = *D_j_* = 0, ∀*j*.

Following van der Merwe (2004), the sigma-points {*v_l_*, 𝓢_*l*_} are defined as

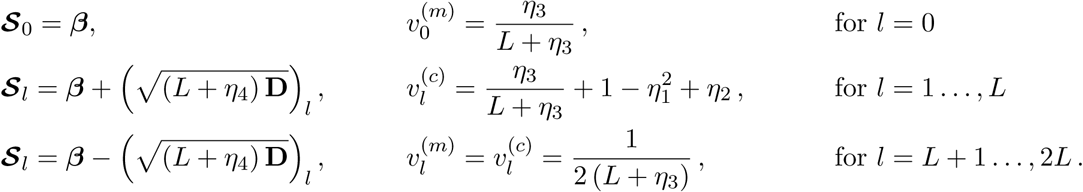

We used *η*_2_ = 2 and *η*_3_ =
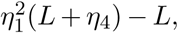
with *η*_1_ = 0.7 and *η*_4_ = 0 as proposed by van der Merwe (2004). The superscripts for *v_l_* indicate whether it is used for the calculation of the mean ^(*m*)^ or the covariance (*c*).

For the examples and applications presented in the manuscript, we assumed that the variability between cells is completely explained by differences in the model parameters. For a set of given parameters, the dynamics of individual cells were described by the RRE (3). Accordingly, the images of the sigma-points in the state and the observation space, 𝓧_*l*_ and 𝓨_*l*_, were computed as

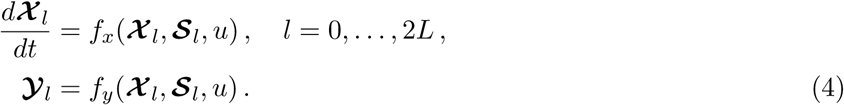

The mean and covariances of the species were computed as

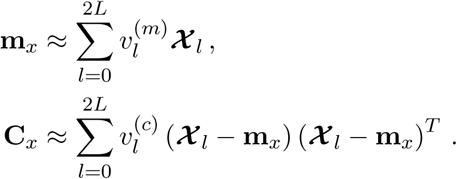

The mean and covariances of the observables read

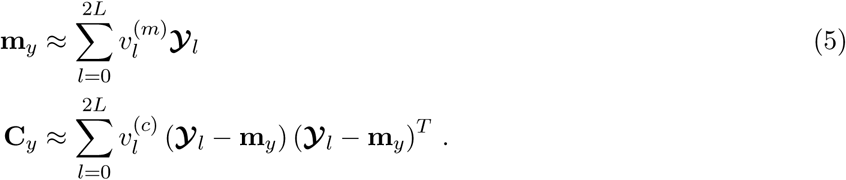

In our MATLAB SPToolbox, the parametrization of **D** was implemented by either a diagonal matrix logarithm or a matrix logarithm (Williams, 1999), in case of correlations between parameters. For our study, we assumed a log-normal distribution of the parameters, i.e., ***β*** and **D** described the median and scale matrix of the corresponding log-normal distribution and the exponent of 𝓢_*l*_ was used in (4).

#### Moment-closure approximation

In this study, we also considered intrinsic variability of biochemical reactions as introduced by discreteness and stochasticity of biochemical reactions. The single-cell dynamics are described by continuous time discrete state Markov chains (CTMCs). We approximated the time-dependent moments of this process using the moment-closure approximation (Engblom, 2006; Lee et al., 2009). This method provided equations for the temporal evolution of moments of the species, i.e., the mean

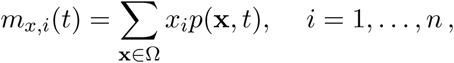

of species *x_i_*, and higher order moments such as the covariance

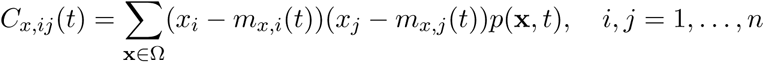

between species *x_i_* and *x_j_*. Here, *p*(*x*, *t*) denotes the chemical master equation, Ω the set of possible states, and *n* the number of species. Given the moments of the species, we calculated the moments of the observables by

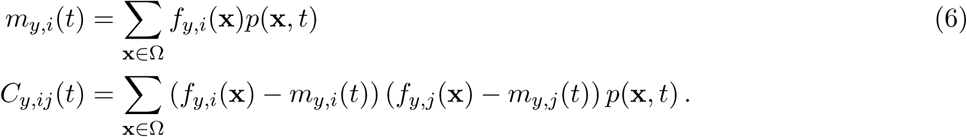

For the automatic generation of the moment-closure approximation and the corresponding simulation files, we employed the MATLAB toolbox CERENA (Kazeroonian et al., 2016). In addition, this toolbox provided the equations for the system size expansion, which can also be incorporated into our modeling framework as an alternative to the moment equations.

### Mechanistic hierarchical population model

For the hierarchical population model, the mechanistic description of individual subpopulations, as introduced in the previous section, is combined with mixture models to describe the entire cell population.

#### Hierarchical model and its approximations

We considered heterogeneous cell populations consisting of multiple subpopulations, *s* = 1,…,*N*. Assuming independence, the distribution of the states and observables in the overall population is the weighted sum of the distribution of the states and observables in the subpopulations, *p_s_*(**x**|*t*) and *p_s_*(**y**|*t*). The weights *w_s_*(*t*) are the relative populations sizes, with ∀*t*: ∑_*s*_ *w_s_*(*t*) = 1. This yields the hierarchical population model

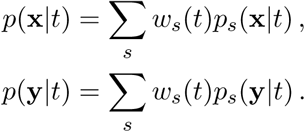

The distribution of states and observables in the subpopulations originate according to the single cell properties. As the measurements ȳ are in general noise corrupted, **ȳ** ~ *p*(ȳ|**y**), we also considered the distribution

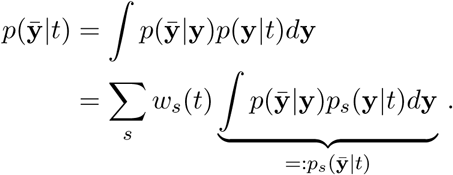

To ensure computational efficiency, the probability distributions *p_s_*(**x**|*t*),*p_s_*(**y**|*t*) and *p_s_*(**ȳ**|*t*) were approximated using the statistical moments. For the measured observables, the computed statistical moments were encoded in ***φ**_s_*, yielding

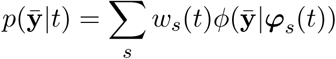

with parametric probability distribution *ϕ*. In this study, we employed the multivariate normal distribution

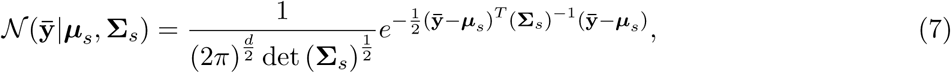

and multivariate log-normal distribution

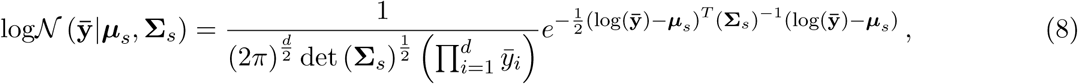

with distribution parameters ***φ**_s_* = (***μ***, **Σ**_*s*_,). For example, for the multivariate normal distribution and no measurement noise, the distributions parameters were obtained by ***μ**_s_* = **m**_*s*,*y*_ and **Σ**_*s*_ = **C**_*s*,*y*_.

#### Likelihood function

The parameters of the hierarchical population model ***θ*** comprise the means/medians of the cell parameters ***β***, ***β**_s_* as well as the entries of the scale matrices **D**, **D**_*s*_, the mixture weights *w_s_*, and measurement noise **Γ**. These parameters were estimated using maximum likelihood estimation. The likelihood function for multivariate measurement data **ȳ**^*e*,*k*,*j*^ ∈ ℝ^*d*^ is given by

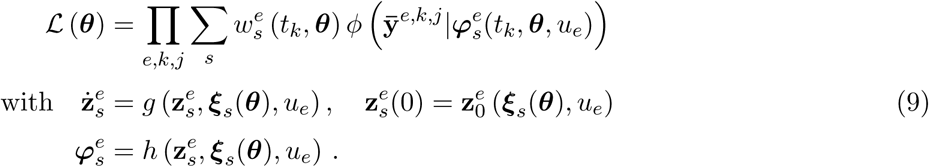

with means and covariances
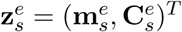
of species **x**. The means and covariances are provided by some map *g*, e.g., the sigma-point approximation or the moment-closure approximation. The subpopulation parameters ***ξ**_s_* = (***β**_s_*, ***D**_s_*) are given by

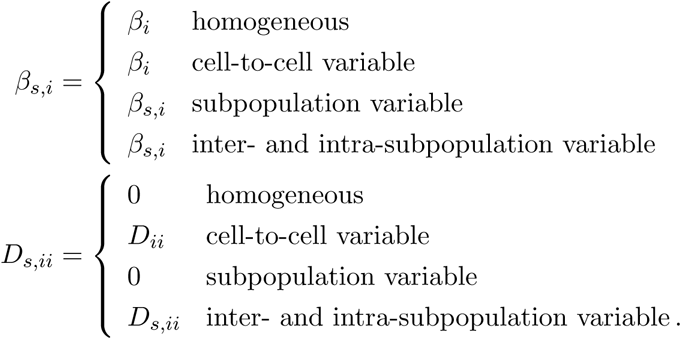

The mapping *h* links the computed moments to the moments of the measurand including measurement noise, which are denoted by **m**_*y*_ = (*m*_*y*, 1_,…, *m*_*y*,*d*_) and **C**_*y*_ and can be calculated as described, e.g., in (5) and (6). For a mixture of normal distributions (7), the means and covariances were linked to the parameters of the normal distribution

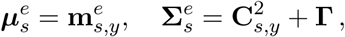

including additive normally distributed measurement noise parametrized by

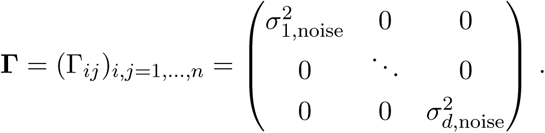

For the log-normal distribution (8), the distribution parameters were directly simulated with the sigma-point approximation for the logarithm of the observable, yielding the relation

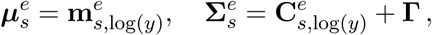

accounting for multiplicative log-normally distributed measurement noise. Alternatively, the mean of the simulation was linked to the mean of the log-normal distribution by

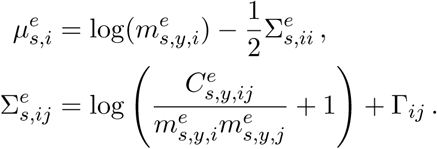

In principle also other distributions can be incorporated in the presented modeling framework. Due numerical reasons, we used the log-likelihood function (Loos et al., 2016).

#### Gradient of likelihood function

To promote efficiency of the numerical optimization and robust convergence, we derived the gradient of the log-likelihood function. For this, the gradient of the corresponding mixture distribution *ϕ* with respect to *θ* was calculated using

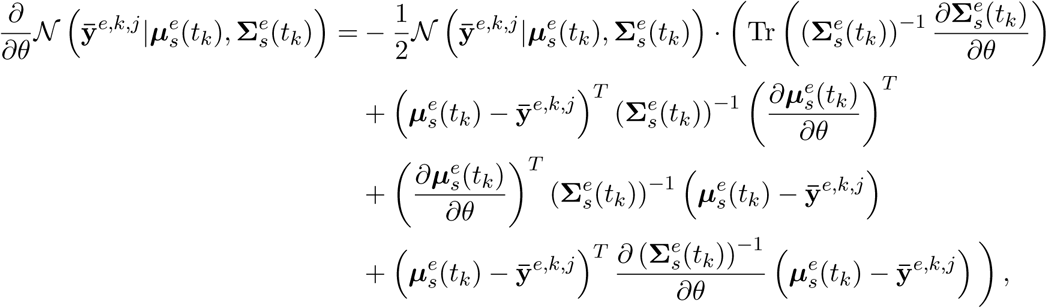

and the relation

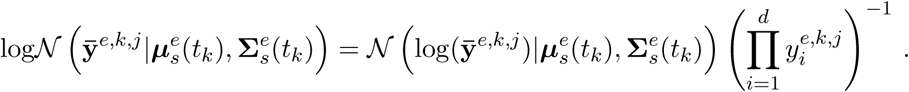

Additionally, the sensitivities of the distribution parameters
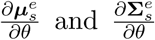
were required, which were obtained by simulating the sensitivity equation for the sigma-point or the moment-closure approximation and mapping it to the distribution parameters using *h*.

### Comparison with existing models

A comparison of the hierarchical population model with existing methods is given in the following:

| method | mechanistic description of |  | subpopulations | multivariate data | reference |
| --- | --- | --- | --- | --- | --- |
|  | dynamics | variability |  |  |  |
| mixture model |  |  | ✓ | ✓ | e.g., Hastie et al. (2009) |
| moment-closure approximation | ✓ | ✓ |  | ✓ | e.g., Zechner et al. (2012) |
| ODE-constrained mixture model | ✓ |  | ✓ |  | Hasenauer et al. (2014) |
| hierarchical population model | ✓ | ✓ | ✓ | ✓ | this manuscript |

### Parameter estimation

For parameter estimation, we used the MATLAB toolbox PESTO, which employs the function `fmincon.m` for local optimization. We used the interior-point algorithm and provided the analytic gradient of the log-likelihood function. Due to numerical better properties, we estimated the log_10_-transformed parameters. To explore the full parameter space, we performed multi-start optimization which has shown to outperform global optimization methods (Raue et al., 2013; Hross and Hasenauer, 2016). For this, randomly drawn initial parameter values were used for the optimization. For the uncertainty analysis, we calculated profile likelihoods (Raue et al., 2009) and the confidence intervals using the corresponding PESTO functions.

### Conversion process

In the manuscript, we considered a model of a conversion process. In the following, we provide a detailed description of the data generation and data analysis. We first introduce the single-cell model of the conversion process. Afterwards, we present the results for the model accounting for the mean, and the hierarchical model accounting for the mean and covariances.

#### Single-cell model

The conversion process is described by the following reactions

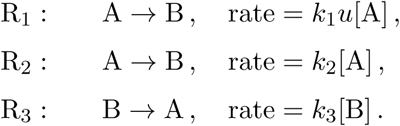

Reaction R_1_ describes the stimulus-dependend conversion, whereas reaction R_2_ models the basal conversion from A to B. The conversion from B to A, reaction R_3_, does not depend on stimulus *u* (Hasenauer et al., 2014). The concentrations of the species A and B are denoted by [A] and [B]. The RRE for (*x*_1_,*x*_2_) = ([A], [B]) is given by

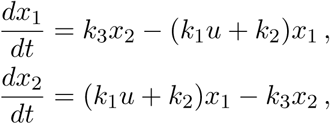

with initial conditions

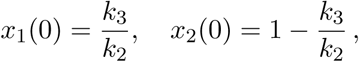

accounting for mass conservation [A] + [B] = 1 and the assumption that the system was in steady state before the stimulus was added at 0 min. We assumed the conversion from B to A to be cell-to-cell variable,

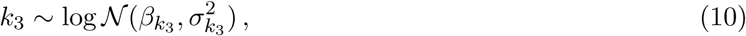

yielding cell-to-cell variable initial conditions. The parameter *k*_1_ was considered to differ between subpopulations and therefore was parametrized by *k*_1,1_ and *k*_1,2_. The weight *w_i_* indicated the proportion of the low responsive subpopulation. We generated artificial data for the parameters

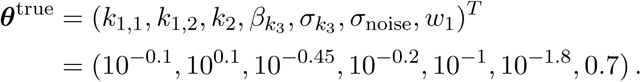

We observed the concentration of B, i.e. *y* = *x*_2_. The data was created including 1000 cells at 5 time points for *u* = 1 by sampling from the distribution for *k*_3_ (10) and simulating the corresponding RREs. Of the 1000 cells, 700 cells belonged to subpopulation 1 with low response to stimulation and 300 cells to the high responsive subpopulation 2. Additionally, the measurements of both subpopulations were assumed to be subject to logarithmic multiplicative measurement noise parameterized by *σ*_noise_. We assumed the parameters ***θ*** to be unknown and estimated them from the data with

i. the approach introduced by Hasenauer et al. (2014) using the means (obtained by the RRE) and
ii. hierarchical population model describing the means and covariances (obtained by the sigma-point approximation)

For both approaches, the underlying subpopulation structure was given, i.e., subpopulation variability of *k*_1_.

#### Hierarchical model using RREs

We considered a hierarchical model with subpopulation means that were described by the RRE. The distribution of the observables was assumed to be log-normal and the scale parameters were estimated from the data. We distinguished the following scenarios:

- one scale parameter that is shared across time points and subpopulations,
- one scale parameter for every subpopulations, which is shared between time points,
- 10 scale parameters that differ for each subpopulation and time-point.

These scale parameters were estimated along with *k*_1,_, *k*_1,2_, *k*_2_,*k*_3_, and *w*_1_ for this setting, which corresponds to the ODE constrained mixture modeling described by Hasenauer et al. (2014). For optimization, the kinetic parameters *k_i_* were assumed to be in the interval [10^−3^,10^3^], the weight *w*_1_ in [0,1], and the scale parameters for the log-normal distribution were restricted to the interval [10^−2^,10^2^]. For each model we performed 50 multi-starts at randomly drawn initial points. The fits corresponding to the optimal parameter values are shown in Figure S1.

#### Hierarchical model using sigma-point approximations

For the hierarchical population model, the parameter vector for subpopulation *s* was given by ***ξ**_s_* = (***β**_s_*, **D**_*s*_) with

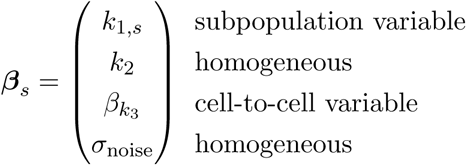

and

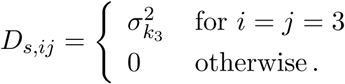

To describe the introduced cell-to-cell variability in *k*_3_ (10) we used the sigma-point approximation for the log-parameters.

To assess whether the true source of heterogeneity can be detected, we tested all possible combinations of additional cell-to-cell variability in *k*_1,*s*_, *k*_2_, or *k*_3_. For this, the sigma-point approximation was applied to the logarithm of the observable, to link the mean and variance of the simulation directly to the distribution parameters of the log-normal distribution. The case of no additional cell-to-cell variability corresponds to the RRE models and is therefore not covered here.

For optimization, the kinetic parameters or their means (in case of cell-to-cell variability) were assumed to be in the interval [10^−3^,10^3^], the scale parameters *σ_k_i__* and measurement noise *σ*_noise_ in [10^−3^, 10^2^] and the weight *w*_1_ in [0,1]. As for the RRE model, we performed 50 multi-starts. The fits corresponding to the optimal parameter values for each model are shown in Figure S1.

To evaluate how the method scales with the number of measured cells, we generated datasets with *n_j_* = {10^1^,10^2^,10^3^, 10^4^,10^5^} measured cells per time point. The average computation time for three replicates for 10 optimization starts for the varying number of data points is shown in Figure S2. The contribution of the evaluation of the density *ϕ*(**ȳ**|*φ_s_*) increased linearly with the number of data points. However, the simulation time was almost constant for increasing number of data points, since the simulation did not depend on the number of measured cells. The slight increase can be explained by the increased number of iterations needed for optimization, which might have occurred due to different effective optimizer tolerances that were not comparable for varying number of data points.

### Differential protein expression

In the main manuscript, we investigated multivariate measurements of differential protein expression. Here, we provide the detailed description of the data generation and the data analysis using the hierarchical model for the full and the marginal distributions.

#### Single-cell model

The simple model of differential protein expression considers six reactions

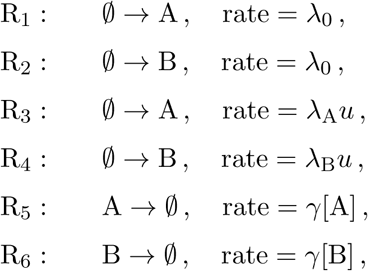

comprising the basal expression with rate λ_0_, degradation with rate *γ* and stimulus-induced expression, depending on *u*, with rate λ_A_ and λ_B_ for protein A and B, respectively. The corresponding ODE system for the temporal evolution of (*x*_1_, *x*_2_) = ([A], [B]) is

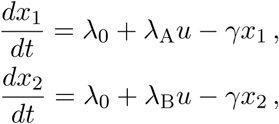

with initial conditions

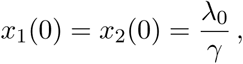

obtained by assuming that the system was in steady state before the stimulus was added at 0 min. Two subpopulations were assumed, one showing high expression of A while the other shows high expression of B after stimulation with *u*. The degradation rate *γ* was considered to be cell-to-cell variable,

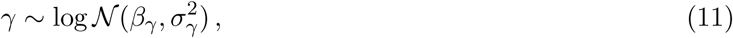

with median *β_γ_* and scale *σ_γ_* which were equal between the subpopulations. The measurements were exposed to log-normally distributed multiplicative measurement noise parametrized by *σ*_noise_.

#### Hierarchical model

The hierarchical model accounted for the subpopulation variability of λ_A_ and λ_B_ and the cell-to-cell variability of *γ*. This yielded the subpopulation parameters

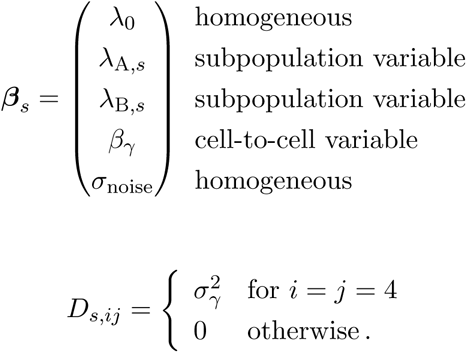

As before, the sigma-point approximation was applied to the log-transformed parameters accounting for the log-normal distribution of *γ*. We performed 100 starts using as data either the full or the marginal distribution of A and B. The parameters and corresponding boundaries are:

| symbol | description | $\theta_{lb}$ | $\theta_{ub}$ | $\theta^{\text{true}}$ |
| --- | --- | --- | --- | --- |
| $\lambda_0$ | basal protein expression | $10^{-3}$ | $10^3$ | $10^{1.7}$ |
| $\lambda_{A,1}$ | induced protein expression of A in subpop. 1 | $10^{-3}$ | $10^3$ | $10^{2.7}$ |
| $\lambda_{A,2}$ | induced protein expression of A in subpop. 2 | $10^{-3}$ | $10^3$ | $10^2$ |
| $\lambda_{B,1}$ | induced protein expression of B in subpop. 1 | $10^{-3}$ | $10^3$ | $10^2$ |
| $\lambda_{B,2}$ | induced protein expression of B in subpop. 2 | $10^{-3}$ | $10^3$ | $10^{2.7}$ |
| $\beta_\gamma$ | median of protein degradation | $10^{-3}$ | $10^3$ | $10^{-1}$ |
| $\sigma_\gamma$ | variability of protein degradation | $10^{-3}$ | $10^1$ | $10^{-1}$ |
| $\sigma_{\text{noise}}$ | measurement noise | $10^{-3}$ | $10^2$ | $10^{-1}$ |
| $w_1$ | weight of subpop. 1 | 0 | 1 | 0.5 |

Using a statistical approach to obtain the number of converged starts (Hross and Hasenauer, 2016), we found that 84/100 starts converged for the full distribution and 91/100 for the marginal distributions.

### NGF-induced Erk1/2 signaling

Here, we provide details for the analysis of NGF-induced Erk1/2 signaling. We employed the model proposed by Hasenauer et al. (2014), which comprises the reactions

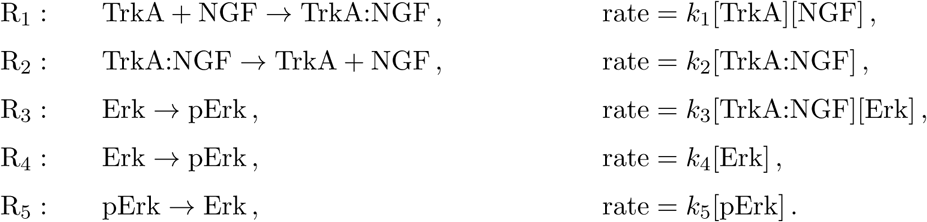

Conservation of mass yields

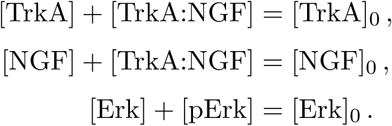

To eliminate structurally non-identifiable parameters, the model was reparametrized to

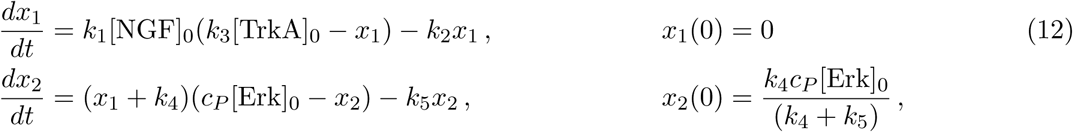

with *x*_1_ = *k*_3_[TrkA:NGF] and *x*_2_ = *c_p_*[pErk]. The observables for the considered experimental conditions are

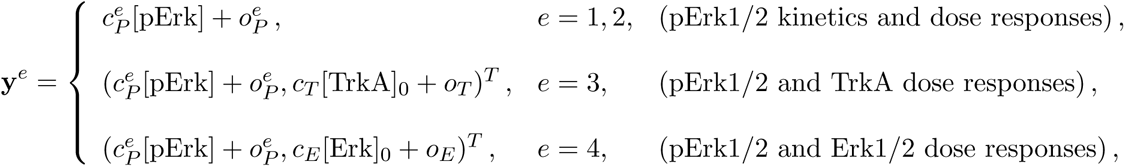

to compare the subpopulations on poly-D-lysine (PDL) and

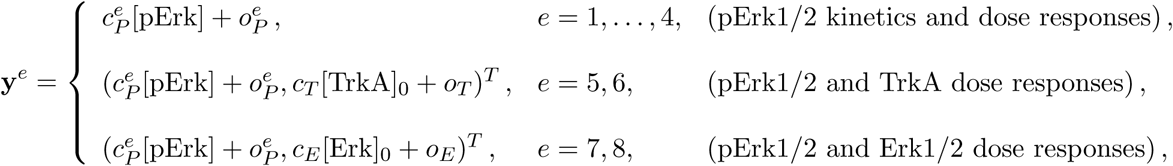

to study the effects of the extracellular scaffolds PDL and collagen type I (Col I) on the neurons (PDL: *e* = 1, 3, 5, 7, Col I: *e* = 2, 4, 6, 8).

The pErk1/2, TrkA and Erk1/2 levels could only be measured up to some scaling constants denoted by *c_p_*, *c_T_* and *c_E_*, respectively, and with some offsets denoted by *o_p_*, *o_T_* and *o_E_*. Each observable was assumed to be subject to multiplicative log-normally distributed measurement noise parameterized by
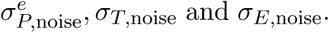
For the comparison of the extracellular scaffold, the same scaling, offset, and measurement noise parameters were used for PDL and Col I. For each subpopulation, we used the sigma-point approximation accounting for cell-to-cell variability in cellular TrkA activity and Erk1/2 levels. The covariance between TrkA activity and relative Erk1/2 expression was parametrized, accounting for correlations, with the matrix logarithm parametrization *M*(*σ_p_*, *σ_p_*, *σ_TE_*) ∈ ℝ^2×2^. All other entries of **D**_*s*_ were assumed to be 0.

#### Data pre-processing

For our analysis, we scaled each replicate such that the quadratic difference of the log-transformed fluorescence mean intensities across replicates is minimal (see `getScalingFactors.m`). The scaled intensities of the cells of each replicate were then pooled and analyzed together.

#### Subpopulation differences

We accounted for all possible combinations of subpopulation variability of *k*_1_, *k*_2_, *k*_4_, *k*_5_, *k*_3_[TrkA]_0_, and
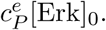
This yielded in total 2^6^ = 64 models that were tested, ranging from *n_θ_* = 26 parameters, for the model assuming no subpopulations at all, to *n_θ_* = 33 parameters, assuming that the subpopulations differ in all parameters. To take into account all hierarchical models, we considered the BIC weights for model *m*

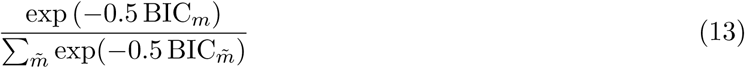

and obtained the BIC weights of a difference by summing over all models that included the corresponding difference.

#### Dephosphorylation rates

To validate, whether the two subpopulations differ in their dephosphorylation/phosphotase activity (parameterized by *k*_5_), we inhibited cells with the Mek-inhibitor U0126 (10*μ*M). NGF binds to the TrkA+ subpopulation and activates pErk1/2 signaling, whereas GDNF binds to the Ret receptor on the opposing subpopulation (TrkA-) and yields pErk1/2 signaling in this neuronal subgroup. Cells were pre-stimulated for 1h with the combined stimuli NGF (20 ng/ml) and GDNF (100 ng/ml) to obtain responses in both subpopulations. We measured pErk1/2 levels to obtain the dynamics of the dephosphorylation as well as TrkA levels to distinguish the two subpopulations. Cells were considered to belong to the TrkA+ subpopulation if their intensity was above 670 and to the TrkA-subpopulation if their intensity was below 630. The measurements were taken at 0,1, 4, 7,10,13,16,19, 22, 25, 28, 31, 34, and 37 min and collected for four replicates.

To obtain the de-phosphorylation rate *k*_5_, we normalized the values of pErk1/2 to 1 at *t* =0 min and 0 at *t*_max_ = 37 min. We fitted an exponential decay

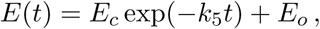

to the scaled data of the four replicates. The scaling *E_c_* and offset *E_o_* could be determined from the boundary conditions

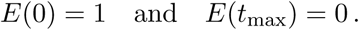

This yielded the four values for the de-phosphorylation in the TrkA+ subpopulation and in the TrkA-subpopulation shown in Figure S3. A two-sample t-test with Welch’s correction gave a p-value of 0.6163, indicating that the dephosphorylation rates in the two subpopulations were not significantly different.

#### Final model

The final model accounted for subpopulation differences in cellular TrkA activity (Figure S4A) and also took into account differences in the variance of TrkA activity between the subpopulations. The fits for the data, which are not shown in the main manuscript are visualized in Figure S4B for the multivariate measurements of pErk1/2 and TrkA, and in Figure 4C for the measurements of pErk1/2 and Erk1/2.

#### Differences mediated by extracellular scaffolds

For the mechanistic comparison of the influence of the extracellular scaffolds, we used the model which assumes subpopulation differences in TrkA levels. The differences between the extracellular scaffolds were parameterized as

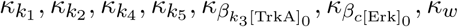

and the parameters were related by

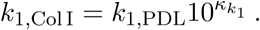

Accounting for these 7 potential differences, we defined 128 hierarchical models. Each model was fitted to the data with multi-start local optimization using at least 20 starts. We sorted the models with respect to their BIC value, for which a low value indicates a good trade-off between model complexity and goodness of fit. The BIC weights for the differences were computed by summing over the BIC weights (13) of the models accounting for the corresponding differences. We found that the best model comprised differences in Erk1/2 expression, Erk1/2 dephosphorylation and cellular TrkA activity. The least suitable model was the model which did not allow differences between the extracellular scaffolds at all. This model was directly followed by the model only accounting for differences in the subpopulation weighting. The fit for the model accounting for differences in Erk1/2 expression, Erk1/2 dephosphorylation and cellular TrkA activity is shown in Figure 6 and Figure S5. The estimated parameters, their boundaries and the 95% confidence interval are

| symbol | description | $\theta_{lb}$ | $\theta_{ub}$ | MLE $\hat{\theta}$ | 95% confidence interval |
| --- | --- | --- | --- | --- | --- |
| $k_1$ | binding affinity | $10^{-6}$ | $10^6$ | 1.514 | [1.114, 2.179] |
| $k_2$ | release of NGF | $10^{-6}$ | $10^6$ | 0.091 | [0.067, 0.131] |
| $k_4$ | basal Erk1/2 phosphorylation | $10^{-6}$ | $10^6$ | 0.014 | [0.012, 0.017] |
| $k_5$ | Erk1/2 de-phosphorylation | $10^{-6}$ | $10^6$ | 0.177 | [0.141, 0.222] |
| $\beta_{k_3[\text{TrkA}]_{0,1}}$ | median of cellular TrkA activity in subpop. 1 | $10^{-6}$ | $10^6$ | $1.9 \cdot 10^{-4}$ | $[1.507 \cdot 10^{-4}, 2.406 \cdot 10^{-4}]$ |
| $\beta_{k_3[\text{TrkA}]_{0,2}}$ | median of cellular TrkA activity in subpop. 2 | $10^{-6}$ | $10^6$ | 0.141 | [0.107, 0.185] |
| $\beta_{c_P^{1,2}[\text{Erk}]_0}$ | median of relative Erk1/2 expression | $10^{-6}$ | $10^6$ | $1.039 \cdot 10^4$ | $[8.989 \cdot 10^3, 1.256 \cdot 10^4]$ |
| $\sigma_{T,1}$ | TrkA variability in subpop. 1 | $10^{-4}$ | $10^4$ | 5.366 | [4.742, 6.121] |
| $\sigma_{T,2}$ | TrkA variability in subpop. 2 | $10^{-4}$ | $10^4$ | 0.303 | [0.277, 0.330] |
| $\sigma_E$ | Erk1/2 variability | $10^{-4}$ | $10^4$ | 0.250 | [0.230, 0.271] |
| $\sigma_{TE}$ | correlation of TrkA and Erk1/2 | $10^{-4}$ | $10^4$ | 2.263 | [2.162, 2.373] |
| $o_P^{1,2}$ | offset pErk1/2 ($e = 1, 2$) | $10^{-6}$ | $10^6$ | $8.064 \cdot 10^{-6}$ | [0, 4.994] |
| $c_T/k_3$ | scaling TrkA | $10^{-6}$ | $10^6$ | $1.735 \cdot 10^4$ | $[1.318 \cdot 10^4, 2.296 \cdot 10^4]$ |
| $o_T$ | offset TrkA | $10^{-6}$ | $10^6$ | 239.7 | [234.3, 245.2] |
| $c_E/c_P^{5,6}$ | scaling Erk1/2 | $10^{-6}$ | $10^6$ | 0.040 | [0.032, 0.049] |
| $o_E$ | offset Erk1/2 | $10^{-6}$ | $10^6$ | 592.2 | [549.8, 631.9] |
| $c_P^{3,4}$ | scaling pErk1/2 ($e = 3, 4$) | $10^{-6}$ | $10^6$ | 0.495 | [0.478, 0.511] |
| $o_P^{3,4}$ | offset pErk1/2 ($e = 3, 4$) | $10^{-6}$ | $10^6$ | 175.4 | [159.8, 190.5] |
| $c_P^{5,6}$ | scaling pErk1/2 ($e = 5, 6$) | $10^{-6}$ | $10^6$ | 0.810 | [0.783, 0.837] |
| $o_P^{5,6}$ | offset pErk1/2 ($e = 5, 6$) | $10^{-6}$ | $10^6$ | 292.1 | [266.2, 317.3] |
| $c_P^{7,8}$ | scaling pErk1/2 ($e = 7, 8$) | $10^{-6}$ | $10^6$ | 1.029 | [0.997, 1.061] |
| $o_P^{7,8}$ | offset pErk1/2 ($e = 7, 8$) | $10^{-6}$ | $10^6$ | 49.91 | [21.91, 76.90] |
| $\sigma_{P,\text{noise}}^{1,2}$ | measurement noise pErk1/2 ($e = 1, 2$) | $10^{-3}$ | $10^1$ | 0.335 | [0.306, 0.361] |
| $\sigma_{P,\text{noise}}^{3,4}$ | measurement noise pErk1/2 ($e = 3, 4$) | $10^{-3}$ | $10^1$ | 0.370 | [0.354, 0.385] |
| $\sigma_{T,\text{noise}}$ | measurement noise TrkA | $10^{-3}$ | $10^1$ | 0.433 | [0.418, 0.448] |
| $\sigma_{P,\text{noise}}^{5,6}$ | measurement noise pErk1/2 ($e = 5, 6$) | $10^{-3}$ | $10^1$ | 0.462 | [0.450, 0.473] |
| $\sigma_{E,\text{noise}}$ | measurement noise Erk1/2 | $10^{-3}$ | $10^1$ | 0.257 | [0.251, 0.263] |
| $\sigma_{P,\text{noise}}^{7,8}$ | measurement noise pErk1/2 ($e = 7, 8$) | $10^{-3}$ | $10^1$ | 0.267 | [0.241, 0.299] |
| $w_1$ | weight of subpopulation 1 | $10^{-4}$ | 1 | 0.294 | [0.289, 0.298] |
| $\kappa_{k_5}$ | difference between extracellular scaffold in $k_5$ | $10^{-3}$ | $10^3$ | 1.257 | [1.221, 1.297] |
| $\kappa_{k_3[\text{TrkA}]_0}$ | difference between extracellular scaffold in TrkA | $10^{-3}$ | $10^3$ | 1.043 | [1.019, 1.068] |
| $\kappa_{c_P^{1,2}\text{Erk}}$ | difference between extracellular scaffold in Erk1/2 | $10^{-3}$ | $10^3$ | 1.393 | [1.357, 1.433] |

### Accounting for intrinsic noise

To study the possibility of accounting for intrinsic noise in the hierarchical population model, we generated artificial data of a two stage gene expression (Figure S6A) using Gillespie’s stochastic simulation algorithm (Gillespie, 1977) incorporated in the MATLAB Toolbox CERENA (Kazeroonian et al., 2016). The system comprises the following reactions

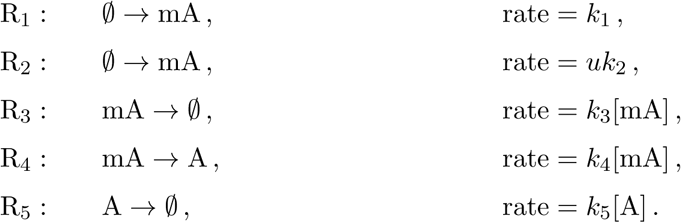

Here, mA denotes the mRNA and A the protein and we assumed that only A could be observed. The two subpopulations differed in their response to stimulus *u* yielding different rates *k*_2,1_ and *k*_2,2_. For this setting, we only accounted for homogeneous and subpopulation variable parameters. However, the intrinsic variability of the births and deaths of individual molecules gave cell-to-cell variability in the cellular states. Cell-to-cell variability in parameters can also be incorporated using the moment-closure approximation.

The ODEs for the temporal evolution of the means and covariances were provided by the toolbox CERENA. In particular, the means *m*_1_ and *m*_2_ and the variances *C*_11_ and *C*_22_ of mRNA mA and protein A, respectively, were described as well as the correlation *C*_12_ of mA and A. The ODE system reads

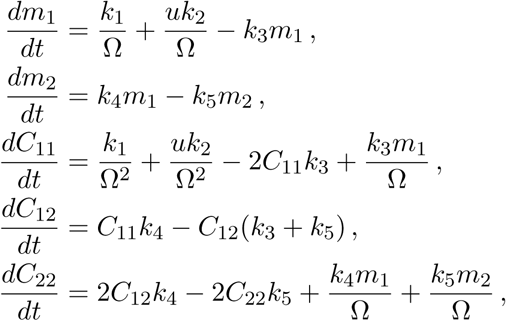

with system size Ω = 1000. Under the assumption that the system was in steady state before stimulation with *u* the initial conditions are

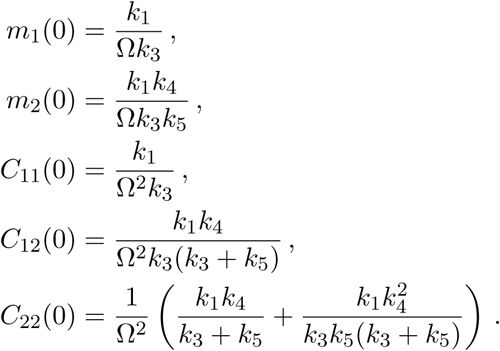

The true parameters used for the generation of the data were

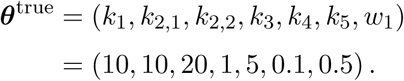

In this example, we employed mixtures of normal distributions, for which the mean and variance were linked to the distribution parameters by ***μ**_s_* = **m**_*s*_ and **Σ**_*s*_ = **C**_*s*_. First, we compared a model accounting for the mean, which was obtained by the RRE (Hasenauer et al., 2014), and a hierarchical model accounting for the mean and covariances, which were obtained by the moment-closure approximation (MA), both accounting for two subpopulations. For the RRE model 10 parameters for the parametrization of the variances were introduced, yielding in total *n_θ_* = 17. The model using the MA only comprised *n_θ_* = 7, since a mechanistic description of the variances was incorporated. For parameter estimation, the kinetic parameters were restricted to the interval [10^−3^,10^3^] and the log_10_-transformed parameters were fitted, whereas the weight *w*_1_ was restricted to [0,1] and fitted linearly. For the RRE model, the parameters for the variance were assumed to lie within [10^−4^,10^2^] and also fitted in log_10_-space. Second, we also studied two models that incorporate the mechanistic description of the variance by the MA, but did not consider the presence of two subpopulations (MA, no subpop.). One of these models, however, accounts for cell-to-cell variability of each parameter (MA, cell-to-cell variability, no subpop.), which corresponds to the description by Zechner et al. (2012).

The models not accounting for subpopulation structures did not fit the data at all (Figure S6B). Even the included variability in parameters did not improve the fit substantially. In contrast, both subpopulation models provided a good fit to the data. However, the BIC for the MA model was substantially better than for the RRE model (BIC_RRE_ – BIC_M_A = 79.09). We found that the MA model gave the optimal value for 40% of the starts and the optimization for the RRE model ended in the optimum for 36% of the starts (Figure S6C). In terms of computation time there was a clear benefit using the mechanistic description of the variance (Figure S6D). The time required for one optimization start was about two-fold faster when using the MA (median = 6.43 sec) instead of RREs (median = 13.13 sec).

Furthermore, we studied the uncertainty of the parameter estimates using profile likelihoods (Figure S6E). Using the MA with subpopulations, all parameters were identifiable, indicated by a narrow profile. This was not the case for RREs, for which some parameters could not be identified from the the data and showed a flat profile. For the case of no subpopulations, most of the true parameters do not lie within the estimated intervals (Figure S6F-G). This emphasizes the importance of taking into account subpopulation structures.

## Statistical analysis

For the analysis of the differences in pErk1/2 activity in the kinetic and dose responses for PDL and Col I, we employed a two-way ANOVA and Sidak’s post-hoc test using GraphPad Prism. For assessing the statistical significance of the predicted differences, we applied the two-sample Welch’s t-test employed by the MATLAB function `ttest2`. Significances are indicated as ^*^ (*p* < 0.05), ^**^ (*p* < 0.01), and ^***^ (*p* < 0.001). Model selection was performed using the BIC. We computed confidence intervals based on the profile likelihoods.

## Data and software availability

The toolbox ODEMM was used to implement the proposed hierarchical modeling framework as well as previous versions. This toolbox also provided the likelihood function and analytical gradient required for parameter estimation. The simulation of the means and covariance using sigma-points was implemented in the SPToolbox. Simulation of the RREs and corresponding sensitivity equations was conducted using the toolbox AMICI (Fröhlich et al., 2016). For the parameter estimation, we employed the toolbox PESTO. All toolboxes and the experimental data are available at https://github.com/ICB-DCM.

**Figure S1:**
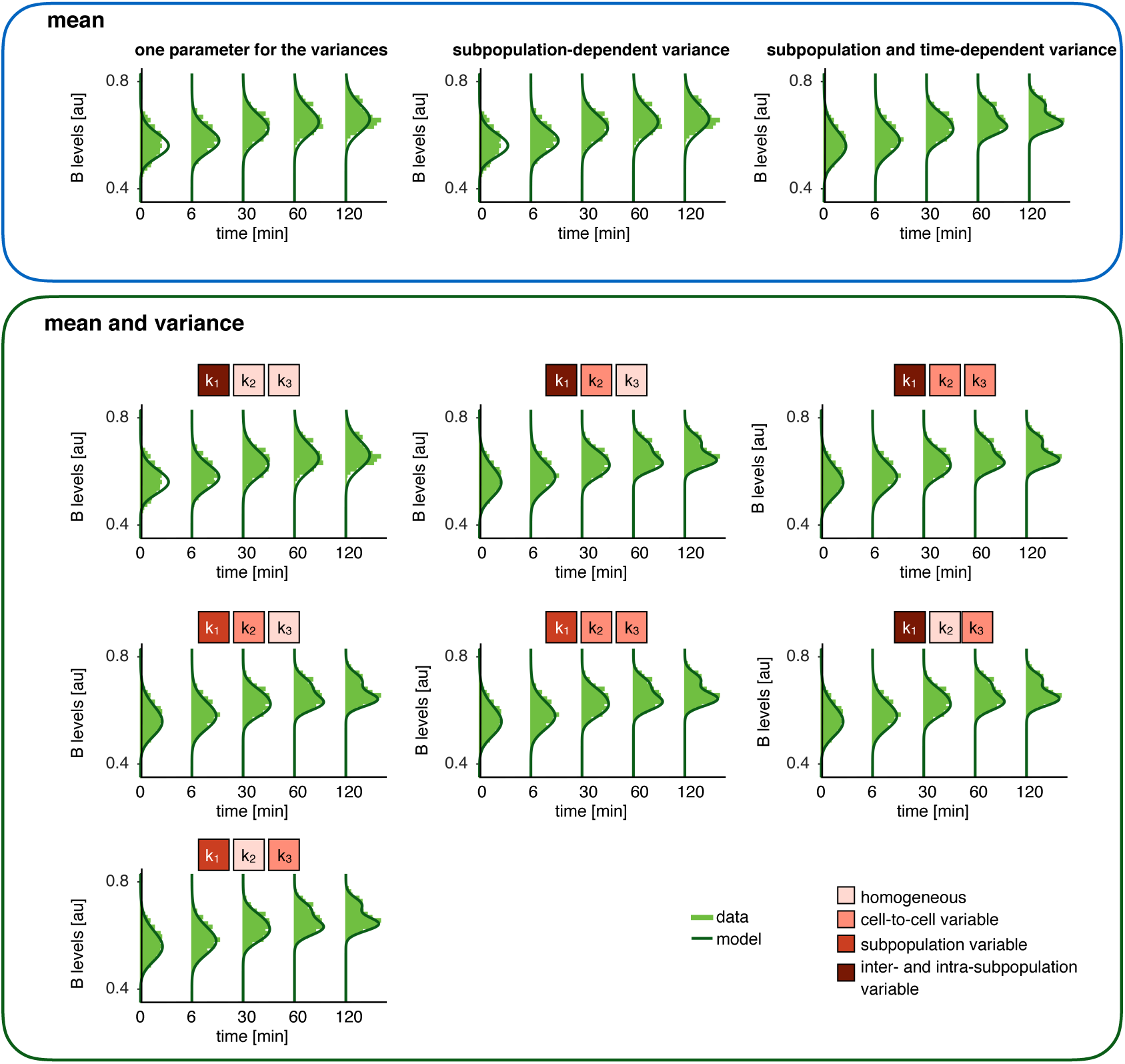
Fitted models for the conversion process. Upper part: Models accounting only for the means using RREs according to (Hasenauer et al., 2014). Lower part: Models accounting for the means and variances using sigma-point approximations.

**Figure S2:**
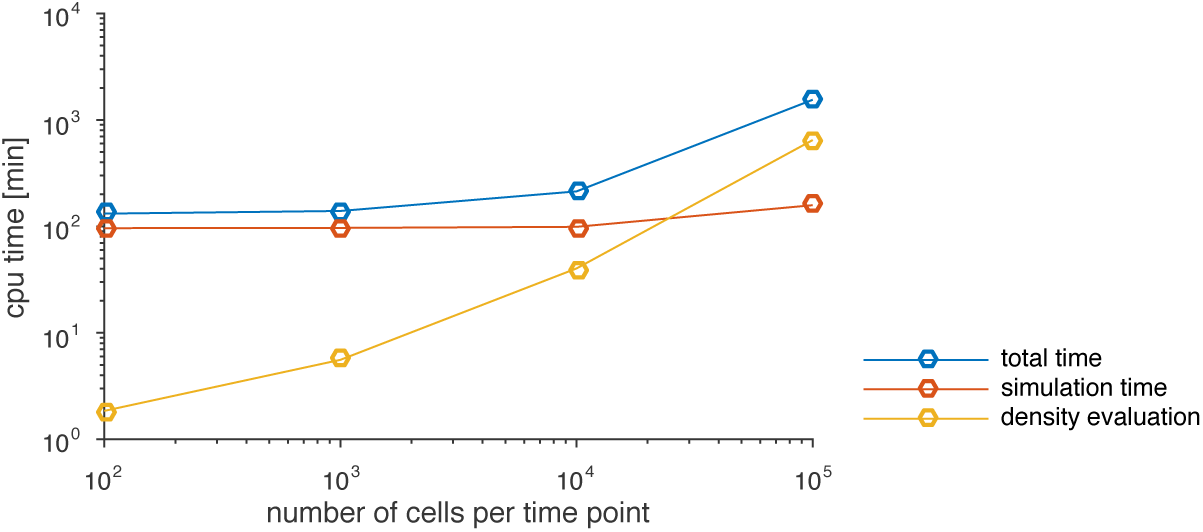
Computation time of the method for the conversion process. The overall computation time for 10 starts is depicted for varying number of measured cells per time point. The circles indicate the mean for three replicates. Different contributions to the overall computation time needed are shown: The time needed for the evaluation of the values for the cells under the density of the mixture distribution (yellow) and the time needed for simulation (orange).

**Figure S3:**
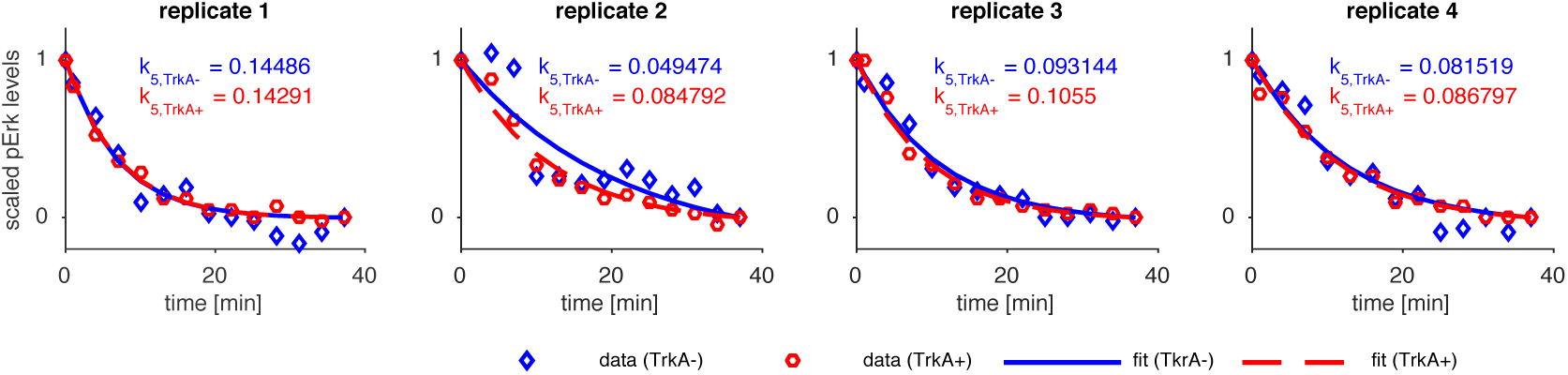
Erk1/2 dephosphorylation in TrkA- and TrkA+ subpopulations. Erk1/2 phosphorylation was induced in both neuronal subgroups TrkA-(blue) and TrkA+ (red) by an 1 h treatment with the combined stimuli NGF (acting on TrkA+ neurons) and GDNF (acting on TrkA-neurons expressing the GDNF receptor Ret). subsequent inhibition of Mek by U0126 induced a pErk1/2 decline. Data of four individual experiments are shown with estimated exponential decay fit. The corresponding values *k*_5_ for the dephosphorylation are noted for both subpopulations.

**Figure S4:**
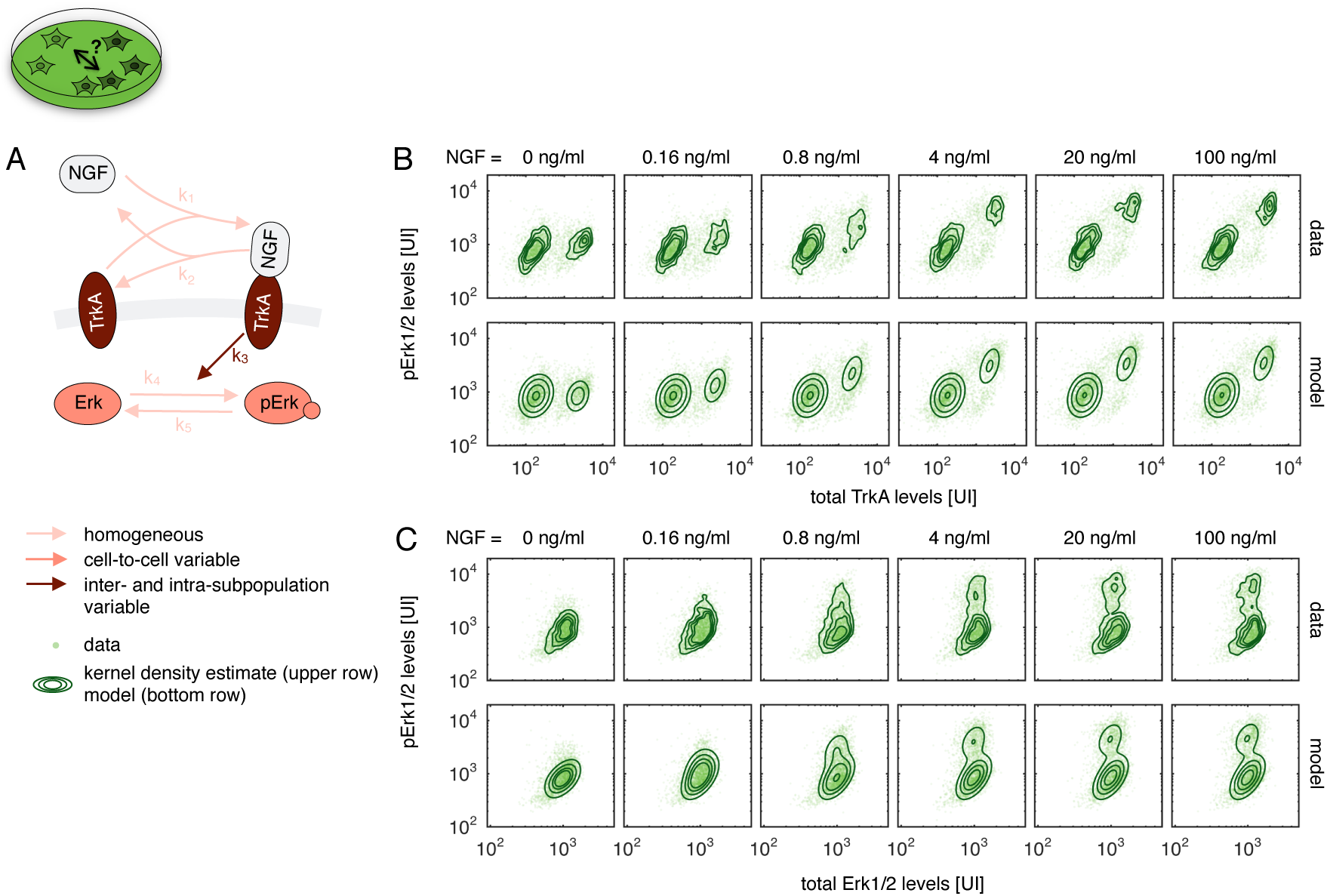
Data and fit for NGF-induced Erk1/2 signaling on PDL. (A) Pathway model with color-coding of the variability of cellular properties. (B,C) Data and fit for combined measurements of (B) TrkA and pErk1/2 levels and (C) Erk1/2 and pErk1/2 levels. The upper rows illustrate the data together with a kernel density estimate. The bottom rows visualize the data together with the contour lines of the hierarchical model.

**Figure S5:**
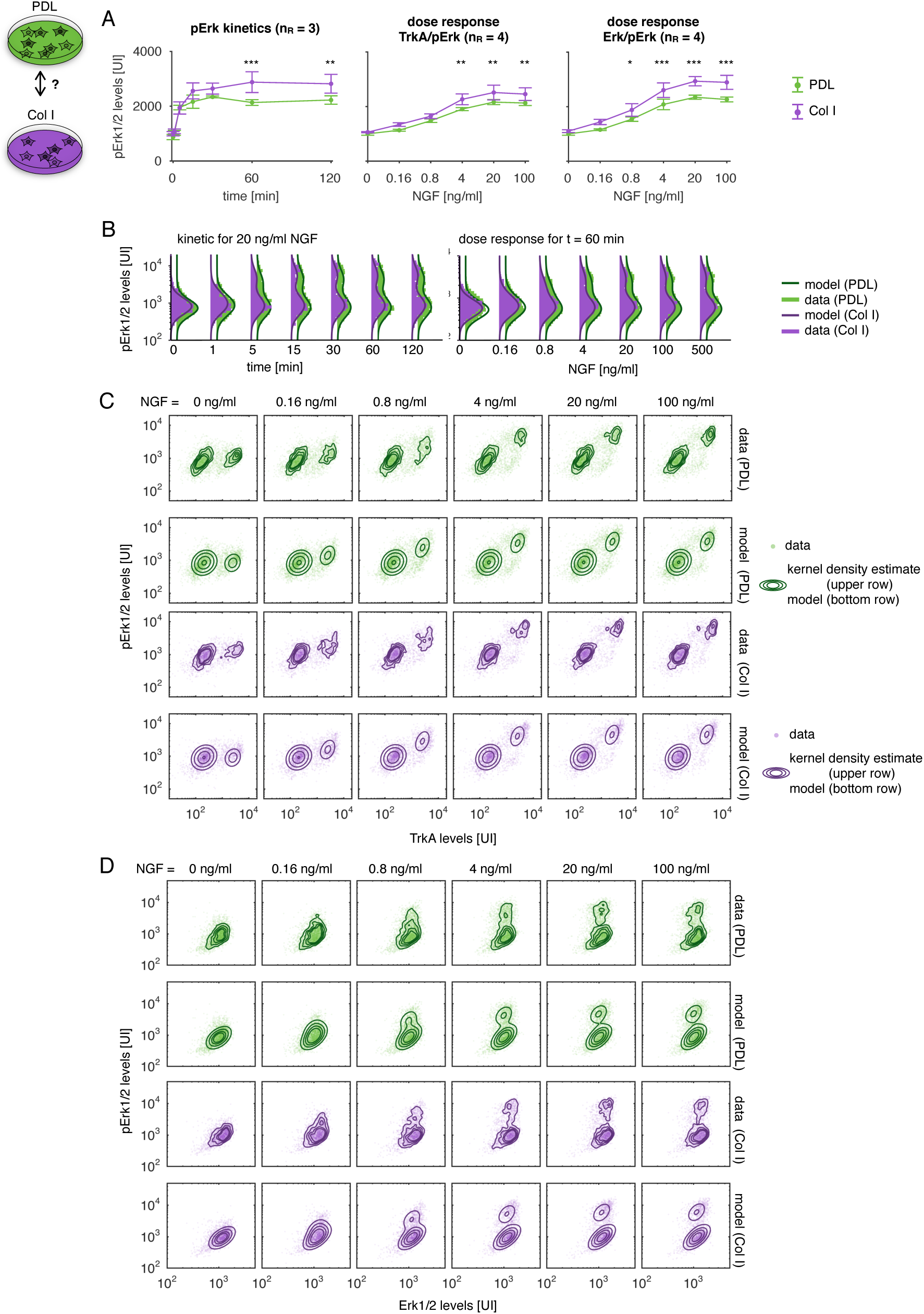
NGF-induced Erk1/2 signaling on different extracellular scaffolds. (A) Mean response to NGF stimulation on Col I compared to PDL. A two-way ANOVA showed significant differences (*p* < 0.001) between the extracellular scaffolds for each experiment. Significances for individual time-points/doses obtained by Sidak’s multiple comparisons test are indicated by ^*^ (*p* < 0.05), ^**^ (*p* < 0.01), and ^***^ (*p* < 0.001). (B-D) Data and fit for NGF-induced Erk1/2 signaling on different extracellular scaffolds. (B) pErk1/2 kinetics and dose responses for PDL (green) and Col I (purple). (C,D) Multivariate measurements of (C) pErk/TrkA levels and (D) pErk/Erk levels. The upper rows illustrate the data together with a kernel density estimate. The bottom rows visualize the data together with the contour lines of the hierarchical model, accounting for differences in Erk1/2 levels, Erk1/2 dephosphorylation, and cellular TrkA activity.

**Figure S6:**
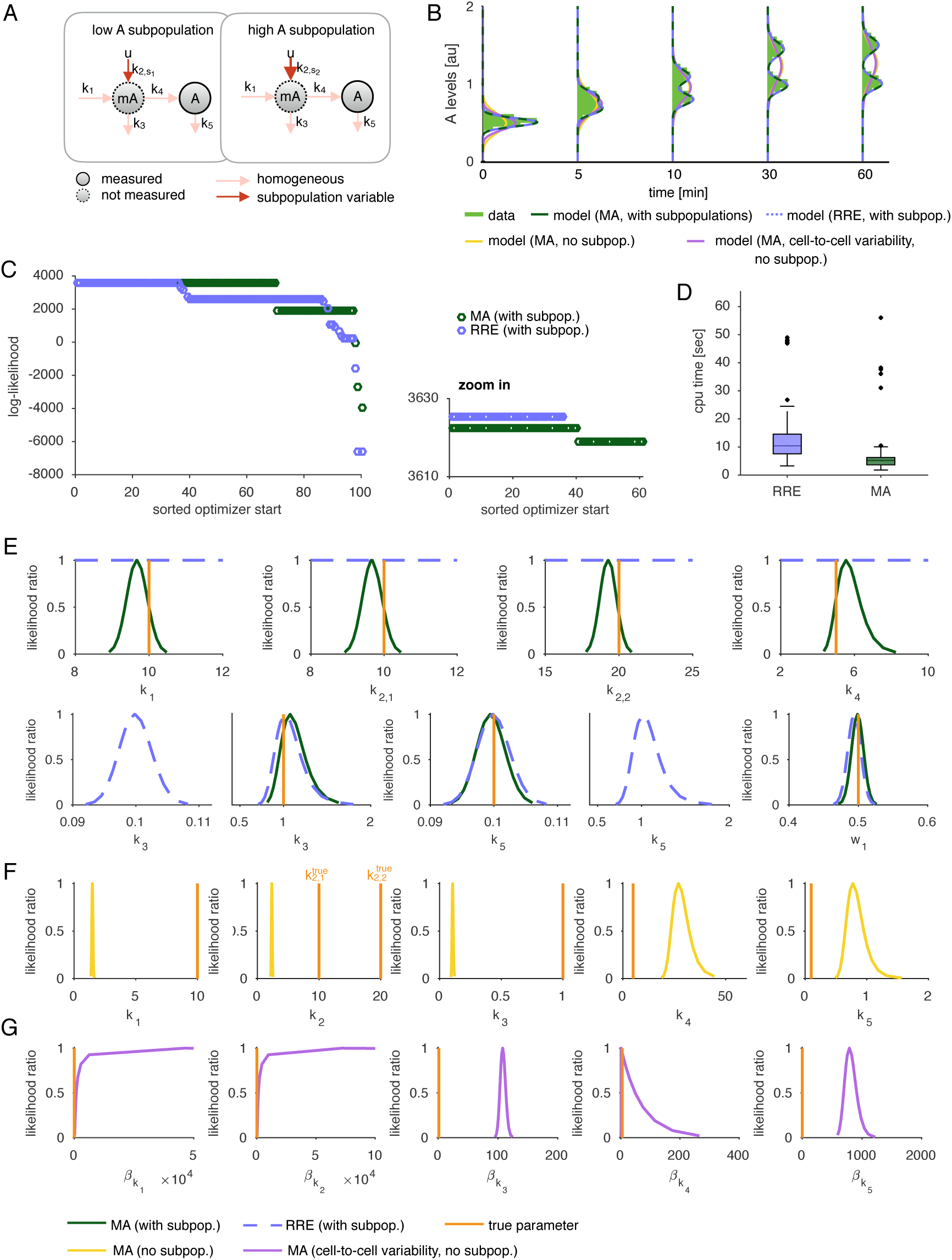
Analysis of a stochastic two stage gene expression model. (A) Illustration of the system. (B) Data and fitted models for the moment-closure approximation (MA), for the case of accounting for subpopulation structures and disregarding subpopulation structures, and reaction rate eqations (RRE). (C) Log-likelihood values for 100 optimization starts sorted decreasingly for MA (green) and RRE (blue). The zoom in shows the best 60 optimization runs. (D) Boxplot for the CPU time needed for one optimizer start. (E) Profile likelihoods of the parameters for the models capturing the subpopulation structure. (F) Profile likelihoods of the parameters for the model using the MA without accounting for subpopulations. (G) Profile likelihoods of the means rates for the model using the MA, accounting for cell-to-cell variability of all parameters but not for subpopulations. This corresponds to the method proposed by Zechner et al. (2012). Note that the range in x-direction differs for subplots (E)-(G).

## References

S. J. Altschuler and L. F. Wu. Cellular heterogeneity: when do differences make a difference? Cell, 141 (4):559–563, 2010

C. Andres, S. Meyer, O. A. Dina, J. D. Levine, and T. Hucho. Quantitative automated microscopy (QuAM) elucidates growth factor specific signalling in pain sensitization. Molecular Pain, 6(98):1–16, 2010.

C. Andres, J. Hasenauer, F. Allgöwer, and T. Hucho. Threshold-free population analysis identifies larger DRG neurons to respond stronger to NGF stimulation. PLoS ONE, 7(3):e34257, 2012.

G. Balázsi, A. van Oudenaarden, and J. J. Collins. Cellular decision making and biological noise: from microbes to mammals. Cell, 144(6):910–925, 2011.

V. R. Buchholz, M. Flossdorf, I. Hensel, L. Kretschmer, B. Weissbrich, P. Gräf, A. Verschoor, M. Schiemann, T. Höfer, and D. H. Busch. Disparate individual fates compose robust CD8+ T cell immunity. Science, 340(6132):630–635, 2013.

F. Buettner, K. N. Natarajan, F. P. Casale, V. Proserpio, A. Scialdone, F. J. Theis, S. A. Teichmann, J. C. Marioni, and O. Stegle. Computational analysis of cell-to-cell heterogeneity in single-cell RNA-sequencing data reveals hidden subpopulations of cells. Nat. Biotechnol., 33(2):155–160, 2015.

L. De Vargas Roditi and M. Claassen. Computational and experimental single cell biology techniques for the definition of cell type heterogeneity, interplay and intracellular dynamics. Curr. Opin. Biotechnol., 34:9–15, 2015.

S. Ebinger, E. Z. Özdemir, C. Ziegenhain, S. Tiedt, C. C. Alves, M. Grunert, M. Dworzak, C. Lutz, V. A. Turati, T. Enver, H.-P. Horny, K. Sotlar, S. Parekh, K. Spiekermann, W. Hiddemann, A. Schepers, B. Polzer, S. Kirsch, M. Hoffmann, B. Knapp, J. Hasenauer, H. Pfeifer, R. Panzer-Grümayer, W. Enard, O. Gires, and I. Jeremias. Characterization of rare, dormant, and therapy-resistant cells in acute lymphoblastic leukemia. Cancer Cell, 30(6):849–862, 2016.

M. B. Elowitz, A. J. Levine, E. D. Siggia, and P. S. Swain. Stochastic gene expression in a single cell. Science, 297(5584):1183–1186, 2002.

W. M. Elsasser. Outline of a theory of cellular heterogeneity. Proc. Natl. Acad. Sci. U S A, 81(16): 5126–5129, 1984

S. Engblom. Computing the moments of high dimensional solutions of the master equation. Appl. Math. Comp., 180:498–515, 2006.

S. Filippi, C. P. Barnes, P. D. Kirk, T. Kudo, K. Kunida, S. S. McMahon, T. Tsuchiya, T. Wada, S. Kuroda, and M. P. Stumpf. Robustness of mek-erk dynamics and origins of cell-to-cell variability in mapk signaling. Cell Reports, 15(11):2524–2535, 2016.

F. Fröhlich, P. Thomas, A. Kazeroonian, F. J. Theis, R. Grima, and J. Hasenauer. Inference for stochastic chemical kinetics using moment equations and system size expansion. PLoS Comput. Biol., 12(7): e1005030, 2016.

D. T. Gillespie. Exact stochastic simulation of coupled chemical reactions. J. Phys. Chem., 81(25): 2340–2361, 1977.

P. J. Green. Reversible jump Markov chain Monte Carlo computation and Bayesian model determination. Biometrika, 82(4):711–732, 1995.

J. Hasenauer, S. Waldherr, M. Doszczak, N. Radde, P. Scheurich, and F. Allgöwer. Identification of models of heterogeneous cell populations from population snapshot data. BMC Bioinformatics, 12 (125), 2011.

J. Hasenauer, C. Hasenauer, T. Hucho, and F. J. Theis. ODE constrained mixture modelling: A method for unraveling subpopulation structures and dynamics. PLoS Comput. Biol., 10(7):e1003686, 2014.

T. Hastie, R. Tibshirani, and J. Friedman. The elements of statistical learning, volume 2. Springer, 2009

S. C. Hicks, M. Teng, and R. A. Irizarry. On the widespread and critical impact of systematic bias and batch effects in single-cell RNA-seq data. bioRxiv, page 025528, 2015

S. Hross and J. Hasenauer. Analysis of CFSE time-series data using division-, age- and label-structured population models. Bioinformatics, 32(15):2321–2329, 2016.

T. Hucho and J. D. Levine. Signaling pathways in sensitization: toward a nociceptor cell biology. Neuron, 55(3):365–376, 2007.

J. Isensee, M. Diskar, S. Waldherr, R. Buschow, J. Hasenauer, A. Prinz, F. Allgöwer, F. W. Herberg, and T. Hucho. Pain modulators regulate the dynamics of PKA-RII phosphorylation in subgroups of sensory neurons. J. Cell Sci., 127:216–229, 2014.

S. Islam, A. Zeisel, S. Joost, G. La Manno, P. Zajac, M. Kasper, P. Lonnerberg, and S. Linnarsson. Quantitative single-cell RNA-seq with unique molecular identifiers. Nat. Methods, 11(2):163–166, 2014.

R. -R. Ji, R. W. Gereau, M. Malcangio, and G. R. Strichartz. MAP kinase and pain. Brain Res Rev., 60 (1):135–148, 2009.

A. Kazeroonian, F. Fröhlich, A. Raue, F. J. Theis, and J. Hasenauer. CERENA: ChEmical REaction Network Analyzer–A toolbox for the simulation and analysis of stochastic chemical kinetics. PLoS One, 11(1):e0146732, 2016.

P. V. Kharchenko, L. Silberstein, and D. T. Scadden. Bayesian approach to single-cell differential expression analysis. Nat. Methods, 11(7):740–742, 2014.

H. Koeppl, C. Zechner, A. Ganguly, S. Pelet, and M. Peter. Accounting for extrinsic variability in the estimation of stochastic rate constants. Int. J. Robust Nonlinear Control, 22(10):1–21, 2012.

C. H. Lee, K. H. Kim, and P. Kim. A moment closure method for stochastic reaction networks. J. Chem. Phys., 130(13):134107, 2009.

G. Lee, W. Finn, and C. Scott. Statistical file matching of flow cytometry data. J. Biomed. Inform., 44 (4):663–676, 2011.

C. Loos, A. Fiedler, and J. Hasenauer. Parameter estimation for reaction rate equation constrained mixture models. In E. Bartocci, P. Lio, and N. Paoletti, editors, Proceedings of the 13th Computational Methods in Systems Biology, Cambridge, UK, volume 9859 of LNCS, pages 186–200. Springer International Publishing, 2016.

C. Maier, C. Loos, and J. Hasenauer. Robust parameter estimation for dynamical systems from outlier-corrupted data. Bioinformatics, 33(5):718–725, 2017.

M. Malik-Hall, O. A. Dina, and J. D. Levine. Primary afferent nociceptor mechanisms mediating ngf-induced mechanical hyperalgesia. Eur. J. Neurosci., 21(12):3387–3394, 2005.

V. Moignard, I. C. Macaulay, G. Swiers, F. Buettner, J. Schütte, F. J. Calero-Nieto, S. Kinston, A. Joshi, R. Hannah, F. J. Theis, S. E. Jacobsen, M. F. de Bruijn, and B. Göttgens. Characterization of transcriptional networks in blood stem and progenitor cells using high-throughput single-cell gene expression analysis. Nat. Cell Biol., 15(4):363–372, 2013.

S. Pyne, X. Hu, K. Wang, E. Rossin, T. Lin, L. Maier, C. Baecher-Allan, G. McLachlan, P. Tamayo, D. Hafler, P. De Jager, and J. Mesirov. Automated high-dimensional flow cytometric data analysis. Proc. Natl. Acad. Sci. USA, 106(21):8519–8124, 2009.

A. E. Raftery. Bayes factors and BIC. Socio. Meth. Res., 27(3):411–417, 1999.

A. Raue, C. Kreutz, T. Maiwald, J. Bachmann, M. Schilling, U. Klingmüller, and J. Timmer. Structural and practical identifiability analysis of partially observed dynamical models by exploiting the profile likelihood. Bioinformatics, 25(25):1923–1929, 2009

A. Raue, M. Schilling, J. Bachmann, A. Matteson, M. Schelke, D. Kaschek, S. Hug, C. Kreutz, B. D. Harms, F. J. Theis, U. Klingmüller, and J. Timmer. Lessons learned from quantitative dynamical modeling in systems biology. PLoS One, 8(9):e74335, 2013.

A. Regev, S. Teichmann, E. S. Lander, I. Amit, C. Benoist, E. Birney, B. Bodenmiller, P. Campbell, P. Carninci, M. Clatworthy, H. Clevers, B. Deplancke, I. Dunham, J. Eberwine, R. Eils, W. Enard, A. Farmer, L. Fugger, B. Gottgens, N. Hacohen, M. Haniffa, M. Hemberg, S. K. Kim, P. Klenerman, A. Kriegstein, E. Lein, S. Linnarsson, J. Lundeberg, P. Majumder, J. Marioni, M. Merad, M. Mh-langa, M. Nawijn, M. Netea, G. Nolan, D. Pe’er, A. Philipakis, C. P. Ponting, S. R. Quake, W. Reik, O. Rozenblatt-Rosen, J. R. Sanes, R. Satija, T. Shumacher, A. K. Shalek, E. Shapiro, P. Sharma, J. Shin, O. Stegle, M. Stratton, M. J. T. Stubbington, A. van Oudenaarden, A. Wagner, F. M. Watt, J. S. Weissman, B. Wold, R. J. Xavier, N. Yosef, and Human Cell Atlas Meeting Participants. The Human Cell Atlas. bioRxiv, page 121202, 2017.

W. Reik. Stability and flexibility of epigenetic gene regulation in mammalian development. Nature, 447 (7143):425–432, 2007.

M. Roederer. Compensation in flow cytometry. Curr. Protoc. Cytom., pages 1–14, 2002

H. Rubin. The significance of biological heterogeneity. Cancer Metastasis Rev., 9(1):1–20, 1990.

G. Sauvageau, P. M. Lansdorp, C. J. Eaves, D. E. Hogge, W. H. Dragowska, D. S. Reid, C. Largman, H. J. Lawrence, and R. K. Humphries. Differential expression of homeobox genes in functionally distinct CD34+ subpopulations of human bone marrow cells. Proc. Natl. Acad. Sci. USA, 91(25):12223–12227, 1994.

D. Schnoerr, G. Sanguinetti, and R. Grima. Validity conditions for moment closure approximations in stochastic chemical kinetics. J. Chem. Phys., 141(8):084103, 2014

T. Schroeder. Long-term single-cell imaging of mammalian stem cells. Nat. Methods, 8(4):30–35, 2011.

S. Tay, J. J. Hughey, T. K. Lee, T. Lipniacki, S. R. Quake, and M. W. Covert. Single-cell NF-κB dynamics reveal digital activation and analogue information processing. Nature, 466:267–271, 2010.

R. van der Merwe. Sigma-point Kalman filters for probabilistic inference in dynamic state-space models. Ph.d. thesis, Oregon Health & Science University, 2004

N. G. van Kampen. Stochastic processes in physics and chemistry. North-Holland, Amsterdam, 3rd edition, 2007

P. M. Williams. Matrix logarithm parametrizations for neural network covariance models. Neural Netw., 12(2):299–308, 1999.

C. Willyard. Cancer therapy: an evolved approach. Nature, 532:166–168, 2016.

J. Yao, A. Pilko, and R. Wollman. Distinct cellular states determine calcium signaling response. Mol. Syst. Biol., 12(12):894, 2016

C. Zechner, J. Ruess, P. Krenn, S. Pelet, M. Peter, J. Lygeros, and H. Koeppl. Moment-based inference predicts bimodality in transient gene expression. Proc. Natl. Acad. Sci. U S A, 109(21):8340–8345, 2012.

Z.-Y. Zhuang, H. Xu, D. E. Clapham, and R.-R. Ji. Phosphatidylinositol 3-kinase activates ERK in primary sensory neurons and mediates inflammatory heat hyperalgesia through TRPV1 sensitization. J. Neurosci., 24(38):8300–8309, 2004.

